# Endothelial FOXC1 and FOXC2 promote intestinal regeneration after ischemia-reperfusion injury

**DOI:** 10.1101/2022.03.03.482713

**Authors:** Can Tan, Pieter R. Norden, Ting Liu, Naoto Ujiie, Xiaocai Yan, Kazushi Aoto, Sagrario Ortega, Isabelle G. De Plaen, Tsutomu Kume

**Affiliations:** Feinberg Cardiovascular and Renal Research Institute, Department of Medicine, Feinberg School of Medicine, Northwestern University, Chicago, Illinois, USA; Department of Pediatrics, Feinberg School of Medicine, Northwestern University, Chicago, Illinois, USA; Department of Biochemistry, Hamamatsu University School of Medicine, Hamamatsu, Japan; Mouse Genome Editing Unit, Biotechnology Program, Spanish National Cancer Research Centre, Madrid, Spain

**Author notes:** Correspondence to: Tsutomu Kume, Feinberg Cardiovascular and Renal Research Institute, Department of Medicine, Northwestern University School of Medicine, 300 E. Superior Street, Chicago, Illinois 60611, USA.

## Abstract

Intestinal ischemia induces mucosal damage while simultaneously activating intestinal stem cells (ISCs), which subsequently regenerate the damaged intestinal epithelium. However, whether paracrine factors secreted from vascular endothelial cells (ECs) - blood and lymphatic ECs (BECs and LECs, respectively) – regulate ISC-mediated regeneration have yet to be elucidated. Here, we identify FOXC1 and FOXC2 as essential regulators of paracrine signaling in regeneration of the small intestine after ischemia-reperfusion (I/R) injury. EC- and LEC-specific deletions of *Foxc1*, *Foxc2*, or both in mice augment I/R-induced intestinal damage by causing defects in vascular regrowth, expression of the chemokine CXCL12 and the Wnt activator R-spondin 3 in BECs and LECs, respectively, and activation of Wnt signaling in ISCs. Treatment with CXCL12 and R-spondin 3 rescues the I/R-induced intestinal damage in EC- and LEC-*Foxc* mutant mice, respectively. This study provides evidence that FOXC1 and FOXC2 are required for intestinal regeneration by stimulating paracrine CXCL12 and Wnt signaling.

## Introduction

Tissue regeneration and repair is essential for maintaining physiological homeostasis and relies on the precise control of molecular networks that regulate, or are regulated by, the vasculature. Endothelial cells (ECs) present in the blood and lymphatic vessels are crucial participants in the vascular-dependent processes that restore damaged tissue because they control the secretion of paracrine factors from both the vessels themselves and nearby cells ^1^. However, the fundamental mechanisms by which the vascular system regulates tissue regeneration and repair are poorly understood. Thus, an adequate understanding of the biological processes that contribute to EC-dependent tissue repair, as well as their roles in the pathogenesis and potential treatment of vascular disease or injury, is crucially dependent on a thorough characterization of how blood/lymphatic vessels control tissue regeneration.

Intestinal ischemia is a life-threatening vascular emergency and can be caused by thrombus formation in the mesenteric vasculature, embolisms that arise as a consequence of cardiopulmonary disease, and disease- or shock-induced declines in perfusion, as well as when blood flow is interrupted by intestinal transplantation ^2, 3^. Moreover, impairments in intestinal microvasculature development contribute to the pathogenesis of neonatal necrotizing enterocolitis (NEC), which is the most common life-threatening gastrointestinal emergency in neonatal patients ^4^. Both homeostasis and repair of the small intestine are mediated via intestinal stem cells (ISCs) ^5, 6^. Active ISCs express the specific marker leucine-rich repeat-containing G protein-coupled receptor 5 (Lgr5) and are located at the base of the crypts of the small intestine where they vigorously proliferate to continuously regenerate the intestinal epithelium. Ischemia/reperfusion (I/R) injury, as well as radiation injury and stresses such as acute inflammation, induce apoptosis in proliferating Lgr5^+^ ISCs ^7, 8^, while ISC regeneration after injury is restored largely by dedifferentiation of crypt cells ^9^. However, the mechanisms that coordinate the role of intestinal ECs in intestinal regeneration and repair, and whether blood and lymphatic ECs (BECs and LECs) in the ISC niche regulate the regenerative activity of ISCs as well as the preservation of the ISC niche after injury, have yet to be fully elucidated.

In adult mice, the proliferation of active ISCs is controlled, in part, by Wnt/β-catenin signaling ^10, 11^, and canonical Wnt/β-catenin signaling is promoted by the cooperative activity of Wnt proteins and R-spondins (RSPO1-RSPO4). Lgr5 functions (with Lgr4 and Lgr6) as a cognate receptor for R-spondins, and R-spondins are expressed in mesenchymal stromal cells of the ISC niche ^12, 13^, including Pdgfra^lo^Grem1+ trophocytes as an essential source of R-spondins (RSPO1- RSPO3, especially RSPO3) ^14^. Foxl1+ mesenchymal cells (telocytes) and other non-epithelial stromal cells express Wnt ligands in the ISC niche ^13, 15, 16^. Both the number of Lgr5^+^ ISCs and the regenerative response to intestinal radiation injury are reduced by cotreatment with R-spondin 2- and R-spondin 3-neutralizing antibodies ^17^, and the mucosal damage induced by intestinal I/R injury can be rescued by treatment with R-spondin 3 ^18^. Most importantly, intestinal RSPO3 ^12, 14, 19^ and Wnt2 ^14^ are highly produced by LECs.

The CXC chemokine CXCL12, also called stromal cell-derived factor 1 (SDF-1), is a homeostatic chemokine expressed in many cell types such as stromal cells, ECs, and fibroblasts in various tissues ^20^. CXCL12 is an essential factor for angiogenesis that involves EC proliferation and migration to form neo-vessel networks ^21^. CXCL12 can be induced by hypoxic stress ^22, 23^ and regulates angiogenesis in an autocrine/paracrine manner by interacting with the CXCR4 and CXCR7 receptors ^24^. In the small intestine, CXCR4 is expressed in the crypt epithelial cells ^25, 26^. CXCR4 deficiency in the intestinal epithelium impairs re-epithelialization after acute dextran sodium sulfate (DSS)-induced injury in mice ^26^, whereas CXCL12- overexpressing mesenchymal stem cells (MSCs), distributed in the villous stroma of the small intestine when administered to mice, activate CXCR4 in the intestinal crypts by activating β- catenin and improve survival and intestinal epithelial recovery after radiation injury ^25^. Thus, CXCL12-CXCR4 signaling and the canonical Wnt/β-catenin pathway appear to interact during intestinal regeneration.

FOXC1 and FOXC2 are closely related members of the FOX transcription factor family and have numerous essential roles in cardiovascular development, health, and disease ^27^. Mutations or changes in the copy number of human *FOXC1* are associated with autosomal-dominant Axenfeld-Rieger syndrome, which is characterized by abnormalities in the eye and extraocular defects ^28^, while inactivating mutations of human *FOXC2* are responsible for the autosomal dominant syndrome Lymphedema-distichiasis, which is characterized by obstructed lymph drainage in the limbs and the growth of extra eyelashes ^29^. There is also some evidence that *Foxc2* haploinsufficiency in mice increases their susceptibility to DDS-induced colitis ^30^; however, the precise function of FOXC1 and FOXC2 in vascular repair and intestinal regeneration after ischemic injury have yet to be determined.

In this study, we report that FOXC1 and FOXC2 in intestinal BECs and LECs contributes to vascular repair and intestinal regeneration after I/R injury by regulating expression of paracrine signaling factors. Inducible, EC- and LEC-specific, single and compound mutant mice for *Foxc1* and *Foxc2* showed that the *Foxc* mutations impair regeneration of the small intestine after I/R injury, accompanied by (1) defective repair of intestinal BECs and LECs, (2) reduced expression of CXCL12 and R-spondin 3 in intestinal BECs and LECs, respectively, and (3) decreased activation of the Wnt/β-catenin pathway in ISCs. Importantly, treatment with either CXCL12 or R-spondin 3 partially rescues the defects in intestinal repair and regeneration associated with EC- and LEC-*Foxc1/c2* deficiency. Together, our data show a new role for FOXC1 and FOXC2 as key mediators of paracrine signaling in the intestinal blood/lymphatic vasculature during post-ischemic intestinal repair/regeneration, and our findings may have important implications for other ischemic conditions that are associated with impairments in tissue regeneration.

## Results

### Foxc1 and Foxc2 expression in the murine intestine

*Foxc1* and *Foxc2* expression in intestinal BECs and LECs was evaluated via quantitative PCR (qPCR) of CD45^−^/CD31^+^ ECs, which include both BECs and LECs isolated from the small intestine in adult mice (Figure 1A). Intestinal ECs are particularly vulnerable to I/R injury, and they undergo apoptosis in response to oxidative stress ^2^. Thus, we next investigated whether I/R injury alters *Foxc1/c2* expression in the small intestine of adult mice. Intestinal I/R injury ^31^ was induced by clamping the superior mesenteric artery for 30 minutes to induce occlusion; then, the clip was removed, and the small intestine was allowed to re-perfuse. *Foxc1/Foxc2* expression was significantly greater in the intestinal CD45^−^/CD31^+^ ECs of mice 4 hours after I/R injury than in the intestinal ECs of sham-operated animals (Figure 1, B and C). These results are consistent with previous studies showing that FOXC2 expression in renal tubular cells of the cortex and outer medulla also increases 24 hours after kidney I/R injury ^32^. Comparison of the relative expression levels of *Foxc1* and *Foxc2* in the intestinal ECs 4 hours after I/R injury (Figure 1D) and with those under sham (Figure 1A) revealed that both *Foxc1* and *Foxc2* were increased to a similar extent after I/R injury. Notably, *Foxc1* was more highly expressed than *Foxc2* under the basal and I/R-induced conditions.

**Figure 1.**
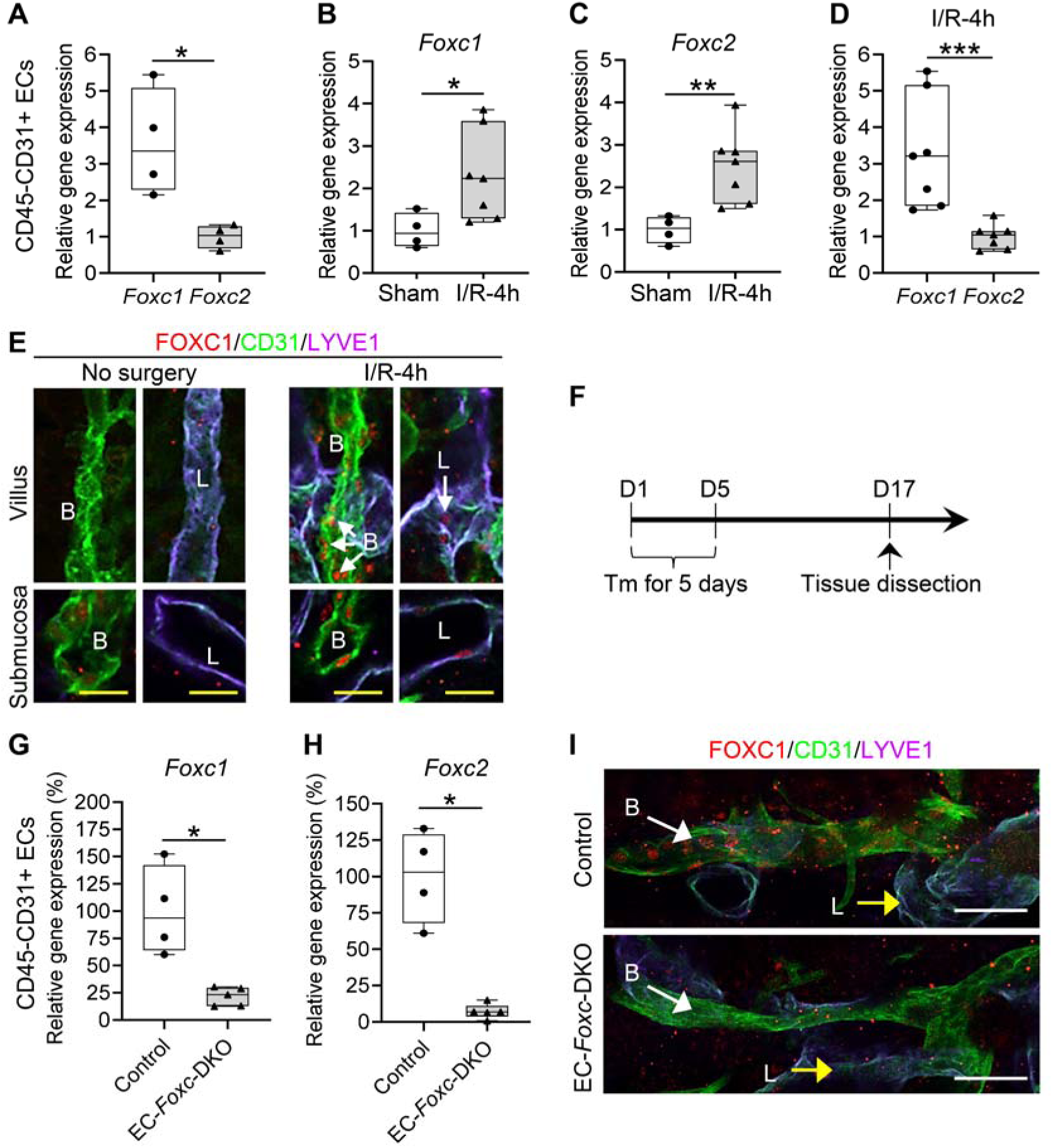
Expression levels of *Foxc1* and *Foxc2* in the mouse small intestine. Relative expression level(s) of **(A)** *Foxc1* and *Foxc2* in sham intestines, **(B)** *Foxc1* and **(C)** *Foxc2* in sham and I/R-4h intestines, **(D)** both *Foxc1* and *Foxc2* at I/R-4h, in the isolated CD45-CD31+ ECs from *Foxc1^fl/fl^;Foxc2^fl/fl^*mouse distal jejuna. Data are box-and-whisker plots, Mann-Whitney *U* test, each symbol represents one mouse, N = 4∼7, \**P*<0.05, \*\**P*<0.01, \*\*\**P*<0.001. **(E)** Representative confocal images of the whole-mount distal jejunum stained with FOXC1/CD31/LYVE1 in *Foxc1^fl/fl^;Foxc2^fl/fl^*mice show the up-regulation of FOXC1 in BECs (B) and LECs (L) at the level of villus and submucosa 4 h after I/R. Scale bars = 20 μm. **(F)** Schematic showing the time of Tamoxifen (Tm) injection and tissue dissection for Figure 1, G-I. Relative expression of *Foxc1* **(G)** and *Foxc2* **(H)** in isolated CD45-CD31+ ECs from distal jejuna 12d after Tm treatment. Data are box-and-whisker plots, Mann-Whitney *U* test, each symbol represents one mouse, N = 4∼5, \**P*<0.05. **(I)** Representative confocal images of submucosal blood vessels (B)/lymphatics (L) in whole-mount intestines stained with FOXC1/CD31/LYVE1. In control, FOXC1 is expressed in the nuclei of the BECs, but hardly detectable in LECs. FOXC1 is down-regulated in BECs after Tm treatment in EC-*Foxc*-DKO mice compared with control. Scale bars = 50 μm. Note that the red tiny spots outside the vasculatures are non-specific staining.

To further investigate FOXC1 expression in the adult mouse intestinal ECs, we performed whole-mount immunostaining labeled with CD31 (EC marker), LYVE1 (LEC marker) and FOXC1. In villus, FOXC1 was hardly detectable in both BECs and LECs, while in submucosa, FOXC1 was weakly expressed in the nuclei of BECs, but was still hardly detectable in submucosal LECs (Figure 1E, no surgery). Four hours after I/R injury, blood and lymphatic vessels were damaged severely, and both FOXC1 and FOXC2 protein levels were upregulated in both intestinal BECs and LECs (Figure 1E and Supplementary Figure 1A). To investigate *Foxc2* expression cells in the intestinal vasculature, we crossed tamoxifen-inducible *Foxc2-Cre^ERT2^* knock-in mice ^33^ with dual Rosa26mTmG reporter mice ^34^. Adult *mTmG/+;Foxc2-Cre^ERT2^* mice were treated with tamoxifen (150 mg/kg) by oral gavage for 5 consecutive days. Consistent with immunohistochemical analysis of FOXC2, *Foxc2*-expressing, recombined EGFP+ cells were also detected in lacteals and villous capillaries of small intestines in adult *mTmG/+;Foxc2- Cre^ERT2^* mice after intestinal I/R injury (Supplementary Figure 1, B and C).

Expression patterns of FOXC1 and FOXC2 in the intestine were also examined in a mouse NEC model ^35^, which includes initial orogastric inoculation of neonatal mice with a standardized adult mouse commensal bacteria preparation and lipopolysaccharide (LPS) to perturb the normal intestinal colonization process, gavage with formula every 3 hours, and exposure to brief episodes of hypoxia for 1 min followed immediately by cold stress (10 min at 4°C) twice daily. With this protocol, about 50-70% of mice typically develop between 36-72 hours intestinal injuries ranging from epithelial injury to transmural necrosis ^35^. At 24 hours after the neonates were subjected to the NEC protocol, immunohistochemical analyses of BEC (CD31 and endomucin [EMCN]) and LEC (PROX1) markers revealed that levels of FOXC1 and FOXC2 were increased in intestinal BECs and LECs (Supplementary Figures 2 and 3), compared to dam- fed littermate controls.

### Generation of tamoxifen-inducible, EC-specific, *Foxc1/c2*-mutant mice in the adult

Murine *Foxc1* and *Foxc2* are both required for vascular development ^36,37,38^, but attempts to determine how the two genes function during pathological (lymph)angiogenesis have been generally unsuccessful, because global single and compound *Foxc1/c2* mutant mice die perinatally with severe cardiovascular abnormalities ^37^. Therefore, we crossed conditional-null *Foxc1^fl^* and *Foxc2^fl^* mutant mice ^39^ with *Cdh5-Cre^ERT2^* mice ^40^ to generate tamoxifen-inducible, EC-specific, compound *Foxc1;Foxc2*-mutant (*Cdh5*-*Cre^ERT2^*;*Foxc1^fl/fl^;Foxc2^fl/fl^*) mice, which (after the mutation is induced) are referred to as EC-*Foxc*-DKO mice ^41^. To induce the mutations, adult mice were treated with tamoxifen (150 mg/kg) by oral gavage for 5 consecutive days, and 12 days after tamoxifen treatment was completed (Figure 1F), qPCR and immunohistochemical analyses confirmed that *Foxc1*- and *Foxc2*-expression was significantly reduced in intestinal ECs of EC-*Foxc*-DKO mice than in the corresponding cells of control littermates (Figure 1, G-H). Importantly, the small intestines of EC-*Foxc*-DKO mice appeared morphologically normal 12 days after tamoxifen treatment (Supplementary Figure 1D), suggesting that EC expression of *Foxc1* and *Foxc2* is not required for maintaining intestinal epithelium homeostasis. We also crossed EC-*Foxc*-DKO mice with Rosa26mTmG mice ^34^, then treated their adult offspring with tamoxifen as described above, and identified recombined EGFP^+^ cells in the small intestines to confirm the efficiency of Cre-mediated recombination in intestinal BECs and LECs (Supplementary Figure 4A).

### EC-specific deletion of *Foxc1* and *Foxc2* decreases intestinal mucosal recovery after I/R

When intestinal I/R injury was induced 12 days after tamoxifen treatment in adult mice (Figure 2A), tissue from the distal jejunum was histologically graded and quantified 24 hours after I/R injury according to the Chiu scoring system ^42^. Compared to the control mice, the intestinal mucosa remained severely injured in EC-*Foxc*-DKO mice (Figure 2, B and C). To further characterize how the loss of EC-specific *Foxc1/c2* expression affects repair of the intestinal mucosa during recovery from I/R injury, we also examined mice carrying tamoxifen-inducible, EC-specific mutations of each individual gene (i.e., EC-*Foxc1*-KO and EC-*Foxc2*-KO mice) and in their control littermates. Recovery from intestinal damage was impaired in both EC-*Foxc1*- KO and EC-*Foxc2*-KO mice 24 hours after I/R injury (Figure 2, D and E; Supplementary Figure 5, A and B). As the extent of intestinal injury in the EC-specific *Foxc* single mutant mice was less than that in the double *Foxc1/c2* mutant mice, EC-*Foxc1* and -*Foxc2* expression is required for intestinal epithelial regeneration. Since I/R-induced local inflammatory response is critically associated with intestinal damage ^18, 43^, mRNA levels of proinflammatory regulators (*TNFα*, *IL-6* and *Cox2*) were measured 3 hours after I/R injury via qPCR. Expression levels of *TNFα* and *Cox2* were significantly higher in EC-*Foxc*-DKO mice than in their control mice (Figure 2F), whereas increased *IL-6* expression in EC-*Foxc*-DKO mice exhibited a trend toward significance (*P*=0.2014). Furthermore, EC-specific loss of *Foxc* genes significantly reduced the proliferative response of intestinal epithelial cells to I/R injury as assessed via immunostaining of EpCAM (intestinal epithelial cell marker) and BrdU (proliferative marker) (Figure 2G), as well as the quantification of BrdU positive epithelial cell number per crypt (Figure 2H).

**Figure 2.**
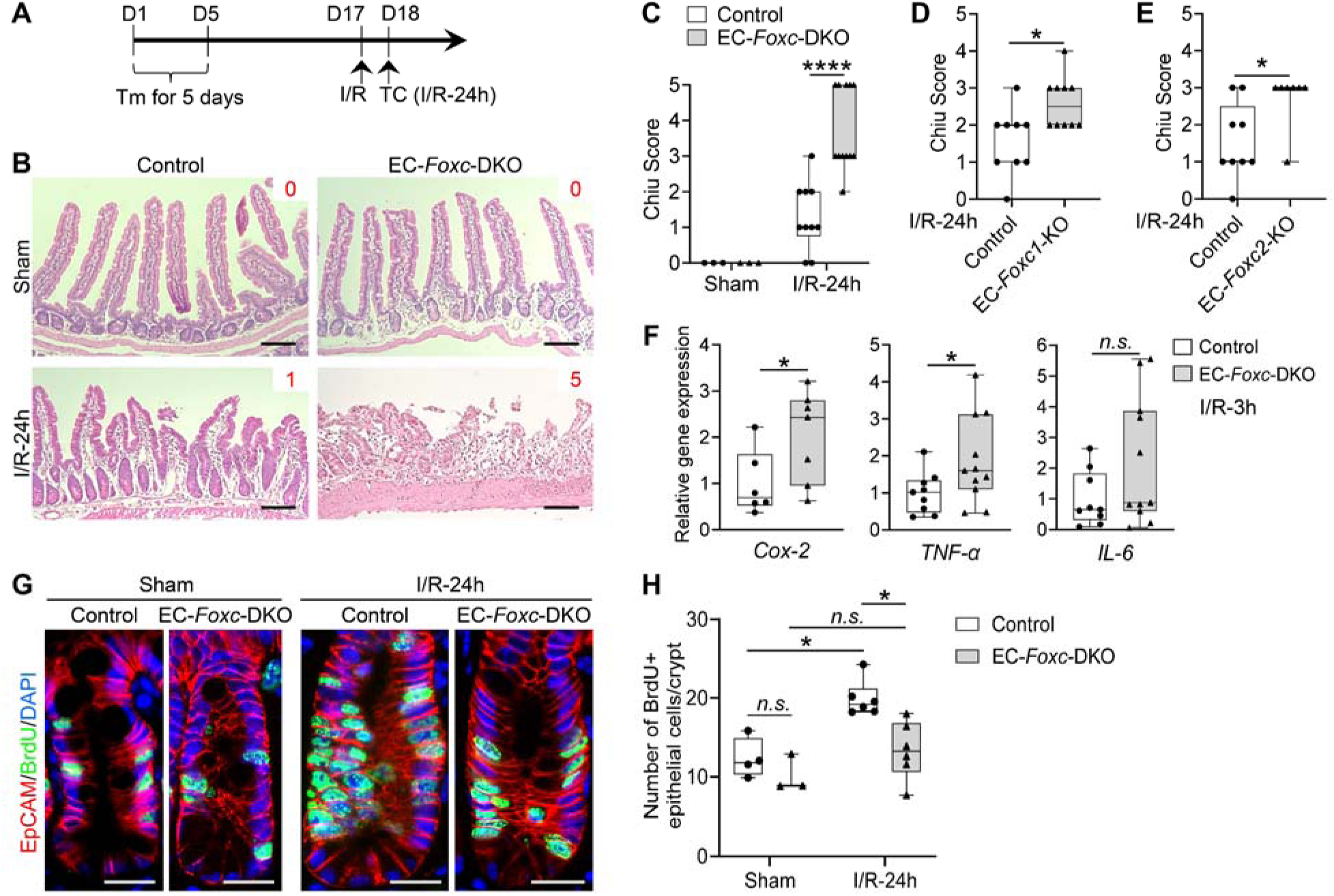
Characterization of defects in the intestinal mucosa in mice with EC-specific deletion of *Foxc1/2* after I/R. (A) Schematic showing the time of Tamoxifen (Tm) injection, I/R surgery and tissue collection (TC). **(B)** Representative H&E staining images of the distal jejuna in EC-*Foxc*-DKO and control mice 24h after I/R. The intestinal ischemic injury grading in the Chiu scoring system is indicated by red numbers (0∼5). Scale bars = 100 μm. Quantification of Chiu Score for **(C)** Control and EC-*Foxc*-DKO, **(D)** Control and EC-*Foxc1*-KO, **(E)** Control and EC-*Foxc2*-KO groups at I/R-24h. Figure 2D and 2E are based on Supplementary Figure 5, A and B. Data are box-and-whisker plots, Mann-Whitney U test, each symbol represents one mouse, N = 3 in C sham groups, N = 7∼12 in C, D, E I/R-24h groups, \**P*<0.05, \*\*\*\**P*<0.0001. **(F)** Relative mRNA expression of proinflammatory mediators *Cox-2*, *TNF-*α and *IL-6* from intestinal tissue lysates at I/R-3h. Data are box-and-whisker plots, Mann-Whitney U test, each symbol represents one mouse, N = 6∼11, \**P*<0.05, *n.s*. = not significant. **(G)** Representative immunostaining images of intestinal crypts labeled with BrdU (proliferative marker, injection performed 2h before tissue collection) and EpCAM (epithelial marker) show the proliferation of epithelial cells in crypts. Paraffin sections (4 µm), scale bars = 20 μm. **(H)** Quantification of the number of BrdU+ epithelial cells per crypt based on Figure 2G. Data are box-and-whisker plots, Kruskal-Wallis One-way ANOVA test, each symbol represents one mouse, N = 3∼6, \**P*<0.05, *n.s*. = not significant.

### Wnt signaling in the small intestine is impaired in EC-*Foxc*-DKO mice after I/R injury

β-catenin regulates the maintenance and regeneration of intestinal epithelial cells ^10, 11^ by translocating from the cytosol to the nucleus of ISCs and altering gene expression in response to activation of the canonical Wnt signaling pathway. This mechanism is consistent with our observations in control mice, because although β-catenin was located at the adherens junctions of epithelial cells in the villus, nuclear β-catenin was detected in ISCs co-immunostained with the ISC marker OLFM4 ^44^ at the crypt base after I/R injury ^13^. However, nuclear localization of β- catenin was impaired (Figure 3, A and B) and the number of cells expressing the Wnt target cyclin D1 (CCND1) ^13^ was significantly reduced in the crypt of EC-*Foxc*-DKO mice both 24 and 48 hours after I/R injury (Figure 3, C and D). Together, the loss of EC-specific *Foxc1/c2* expression impedes I/R-induced Wnt signaling in ISCs.

**Figure 3.**
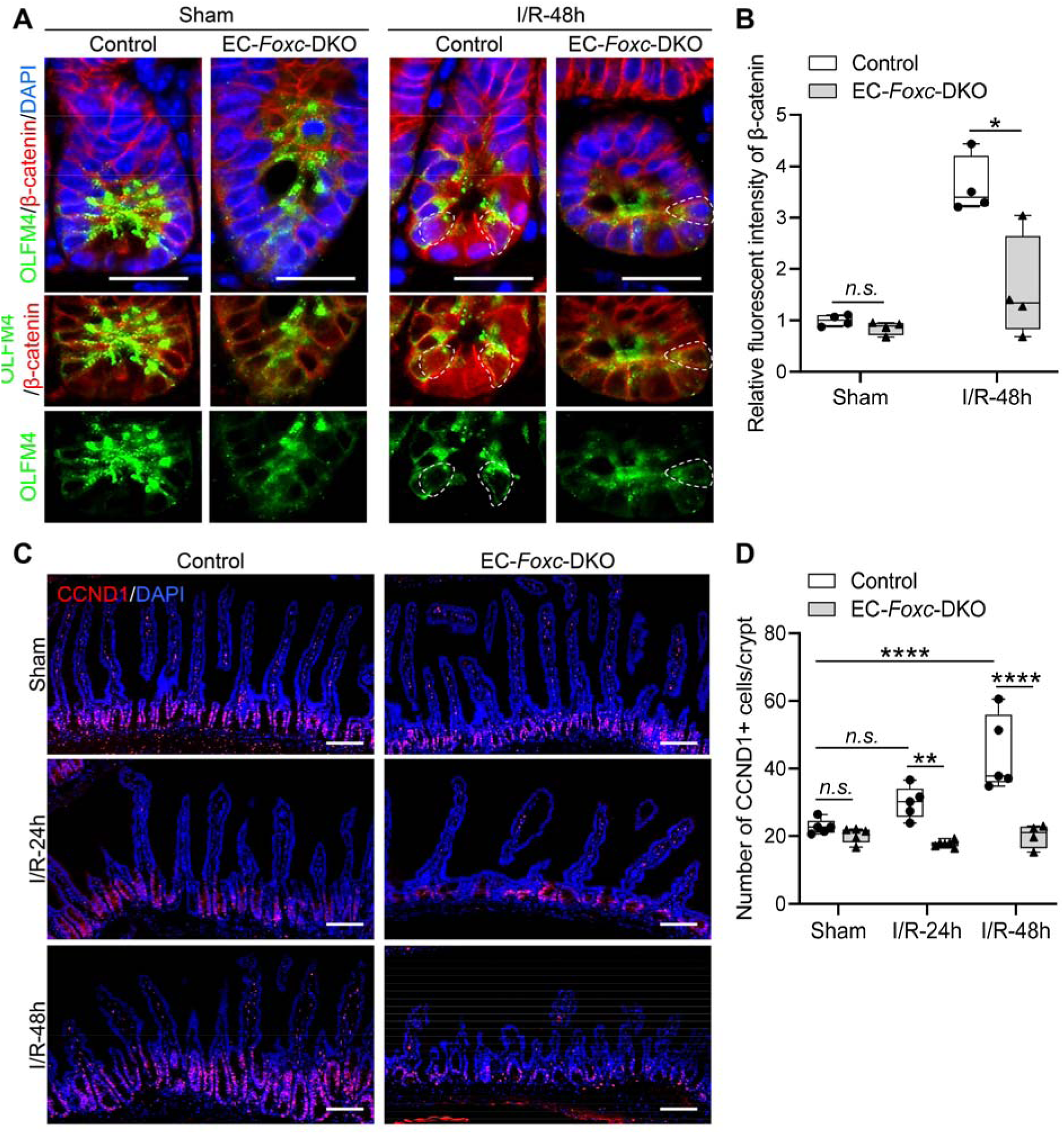
Wnt signaling in the small intestine is impaired in EC-*Foxc*-DKO mice after I/R injury. (A) Representative images of intestinal crypts immunostained with β-catenin and the intestinal epithelial stem cell (ISC) marker OLFM4. At I/R-48h the nuclear translocation of β-catenin in ISCs (dotted circles) was found in control, whereas it’s seldom found in EC-*Foxc*-DKO. Paraffin sections (4 µm), scale bars = 20 μm. **(B)** Quantification of relative fluorescent intensity of β-catenin immunostaining within ISC based on Figure 3A. Data are box-and-whisker plots, Mann-Whitney U test, each symbol represents one mouse, N = 4, \**P*<0.05. **(C)** Representative images of intestinal mucosa labeled with Cyclin D1 (CCND1) in EC-*Foxc*-DKO mice compared with Control group in sham, 24h and 48h after I/R. Paraffin sections (4 µm), scale bars = 100 μm. **(D)** Quantification of the number of CCND1+ epithelial cells per crypt based on Figure 3C. Data are box-and-whisker plots, Kruskal-Wallis One-way ANOVA test, each symbol represents one mouse, N = 4∼6. \*\**P*<0.01, \*\*\*\**P*<0.0001, *n.s*.=not significant.

### Defective mucosal recovery in LEC-specific *Foxc* mutant mice after intestinal I/R injury

As *Cdh5* is expressed in both the blood and lymphatic vasculatures, we generated mice carrying a tamoxifen-inducible, LEC-specific, compound homozygous *Foxc1*^-/-^;*Foxc2*^-/-^ mutation (referred to herein as LEC-*Foxc*-DKO mice) by breeding conditional-null *Foxc1^fl/fl^* and *Foxc2^fl/fl^*mice ^39^ with LEC-specific *Vegfr3-Cre^ERT2^* mice ^45^ to investigate intestinal LEC-specific functions of *Foxc1* and *Foxc2* during I/R. We then confirmed *Vegfr3-Cre*-mediated recombination limited to intestinal LECs of adult LEC-*Foxc*-DKO mice crossed with the Rosa26mTmG reporter mice (Supplementary Figure 4B). The LEC-*Foxc*-DKO mice and their littermate control mice were subjected to the sham or I/R injury procedures, and LEC-specific deletion of *Foxc1* and *Foxc2* significantly increased the severity of intestinal mucosa injury 24 hours after I/R (Figure 4A and Supplementary Figure 5C). Furthermore, increased intestinal damage was also noted in single LEC-*Foxc1*-KO and LEC-*Foxc2*-KO mice after I/R (Figure 4, B and C; Supplementary Figure 5, D and E), suggesting that both FOXC1 and FOXC2 are required in intestinal LECs for intestinal repair in response to I/R injury. In addition, proliferating (BrdU+) intestinal epithelial cells were quantified in the crypts of the LEC-*Foxc*-DKO mice 24 hours after I/R injury, and the LEC-specific loss of *Foxc1/c2* reduced the proliferative response of intestinal epithelial cells to I/R injury (Figure 4D). Consistent with the increased severity of intestinal mucosa defects in the LEC-*Foxc*-DKO mice (Figure 4A), the activation of Wnt signaling (i.e., nuclear localization of β-catenin, Figure 4, E and F) in ISCs and the number of Cyclin D1+ (CCND1+) cells per crypt after intestinal I/R injury were both reduced in the LEC-*Foxc*-DKO mice compared to the control mice (Figure 4G, and Supplementary Figure 6).

**Figure 4.**
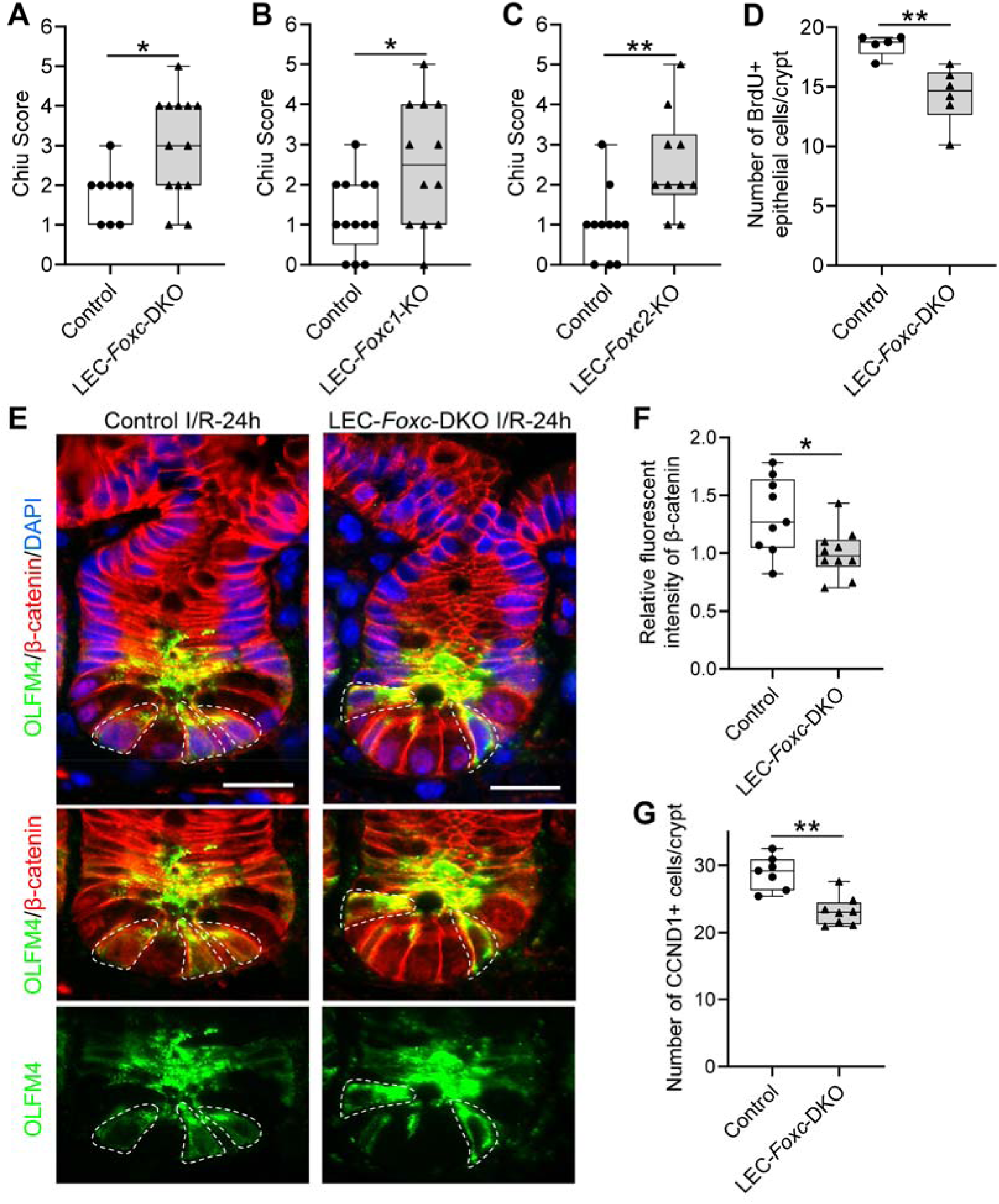
Characterization of defects in the intestinal mucosa in mice with LEC-specific deletion of *Foxc1/2* after I/R. Chiu Score analysis from H&E stained distal jejunum 24h after I/R for: **(A)** Control (*Foxc1^f/f^;Foxc2^f/f^*) and LEC-*Foxc*-DKO (*Vegfr3-Cre^ERT2^;Foxc1^f/f^;Foxc2^f/f^*), **(B)** Control (*Foxc1^f/f^*) and LEC-*Foxc1*-KO (*Vegfr3-Cre^ERT2^;Foxc1^f/f^*), **(C)** Control (*Foxc2^f/f^*) and LEC-*Foxc2*-KO (*Vegfr3-Cre^ERT2^;Foxc2^f/f^*) based on Supplementary Figure 5 C-E. Data are box-and-whisker plots, Mann-Whitney U test, each symbol represents one mouse, N=9∼13, \**P*<0.05, \*\**P*<0.01. **(D)** Quantification of the number of BrdU+ epithelial cells per crypt in control and LEC-*Foxc*-DKO mice at 24h after I/R. Data are box-and-whisker plots, Mann-Whitney U test, each symbol represents one mouse, N = 5∼6. \*\**P*<0.01. **(E)** Representative images of crypts immunostained with OLFM4 and β-catenin in control and LEC-*Foxc*-DKO mice 24h after I/R. The accumulation of β-catenin in the nuclei of ISCs (dotted circles) was found in Control but inhibited in LEC-Foxc-DKO mice. Paraffin sections (4 µm), scale bars = 20 μm. **(F)** Quantification of relative fluorescent intensity of β-catenin immunostaining within ISC based on Figure 4E. Data are box-and-whisker plots, Mann-Whitney U test, each symbol represents one mouse, N = 9∼10, \**P*<0.05. **(G)** Quantification of the number of CCND1+ epithelial cells per crypt at I/R-24h from the intestinal immunostaining of CCND1 based on Supplementary Figure 6. Data are box-and-whisker plots, Mann-Whitney U test, each symbol represents one mouse, N = 7∼8. \*\**P*<0.01.

### Defective vascular recovery in EC- and LEC-specific deletions of *Foxc1/c2* genes after intestinal I/R injury

The vascular recovery of villus BECs and LECs after I/R injury proceeds via a stepwise process in which blood capillaries (BECs) regrow earlier than lacteals (LECs) in the villous stroma ^46^. We found that the EC-*Foxc*-DKO mutation was associated with defective vascular repair of intestinal BECs and LECs after I/R injury (Figure 5). Vascular endothelial growth factor (VEGF) receptor (R) 2 (VEGFR2) and VEGFR3 were highly expressed in the growing tips of villous BECs and LECs, respectively, from control mice after intestinal I/R injury but were severely diminished in EC-*Foxc*-DKO mice (arrows in Figure 5). Notably, proliferation of intestinal BECs and LECs was also significantly reduced (Figure 6, A and B), while the number of apoptotic BECs and LECs was significantly increased (Figure 6, C and D), in EC-*Foxc*-DKO mice than in their control mice after I/R injury. Following whole-mount immunostaining for CD31 and LYVE1 (Supplementary Figure 7, A and B; Supplementary videos), the length of blood capillaries and lacteals (Figure 6E) as well as the percentage of lacteal length to blood capillary length (Figure 6F) were measured. The regrowth of both blood capillaries and lacteals in the villi were significantly decreased in EC-*Foxc*-DKO mice after intestinal I/R injury, whereas EC-*Foxc*-double mutant lacteals were shorter than controls in sham treatments.

**Figure 5.**
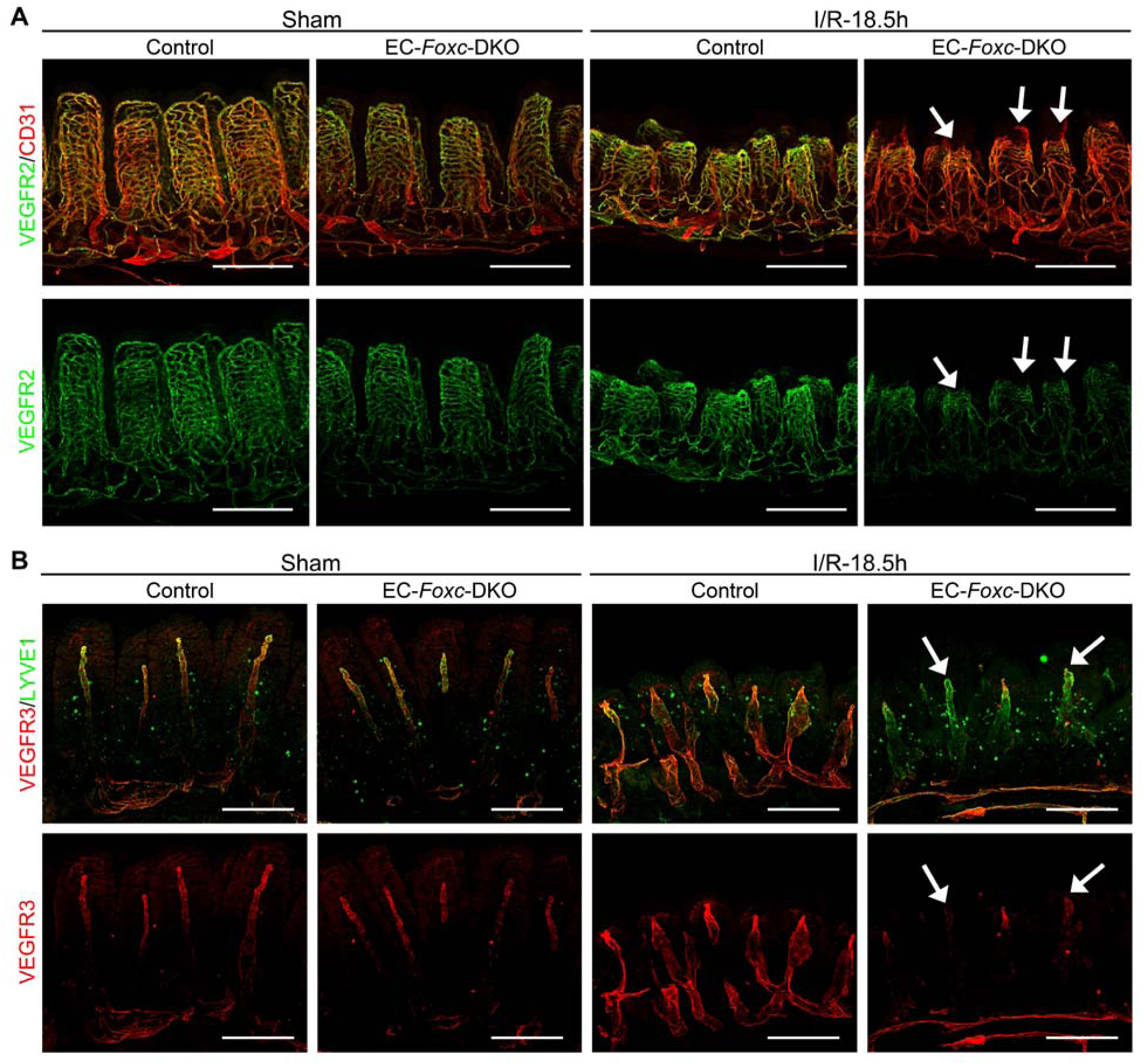
Defects in vascular regeneration after intestinal I/R injury in EC-*Foxc*-DKO mouse. (A) Representative images of intestinal whole-mount VEGFR2/CD31 (green/red) immunostaining show increased VEGFR2 (green) expression at the angiogenic front of villous blood capillaries in control mice at I/R-18.5h. In EC-*Foxc*-DKO, the increase of VEGFR2 is inhibited in villous blood vessels and the repair of blood vasculatures is also impaired (arrow). Scale bars = 200 μm. **(B)** Representative images of intestinal whole-mount VEGFR3/LYVE1 (red/green) immunostaining. 18.5h after I/R, VEGFR3 (red) is increased in lacteals especially at the lacteal tips in the control but is inhibited in the EC-*Foxc*-DKO lacteals (arrow). Scale bars = 200 μm.

**Figure 6.**
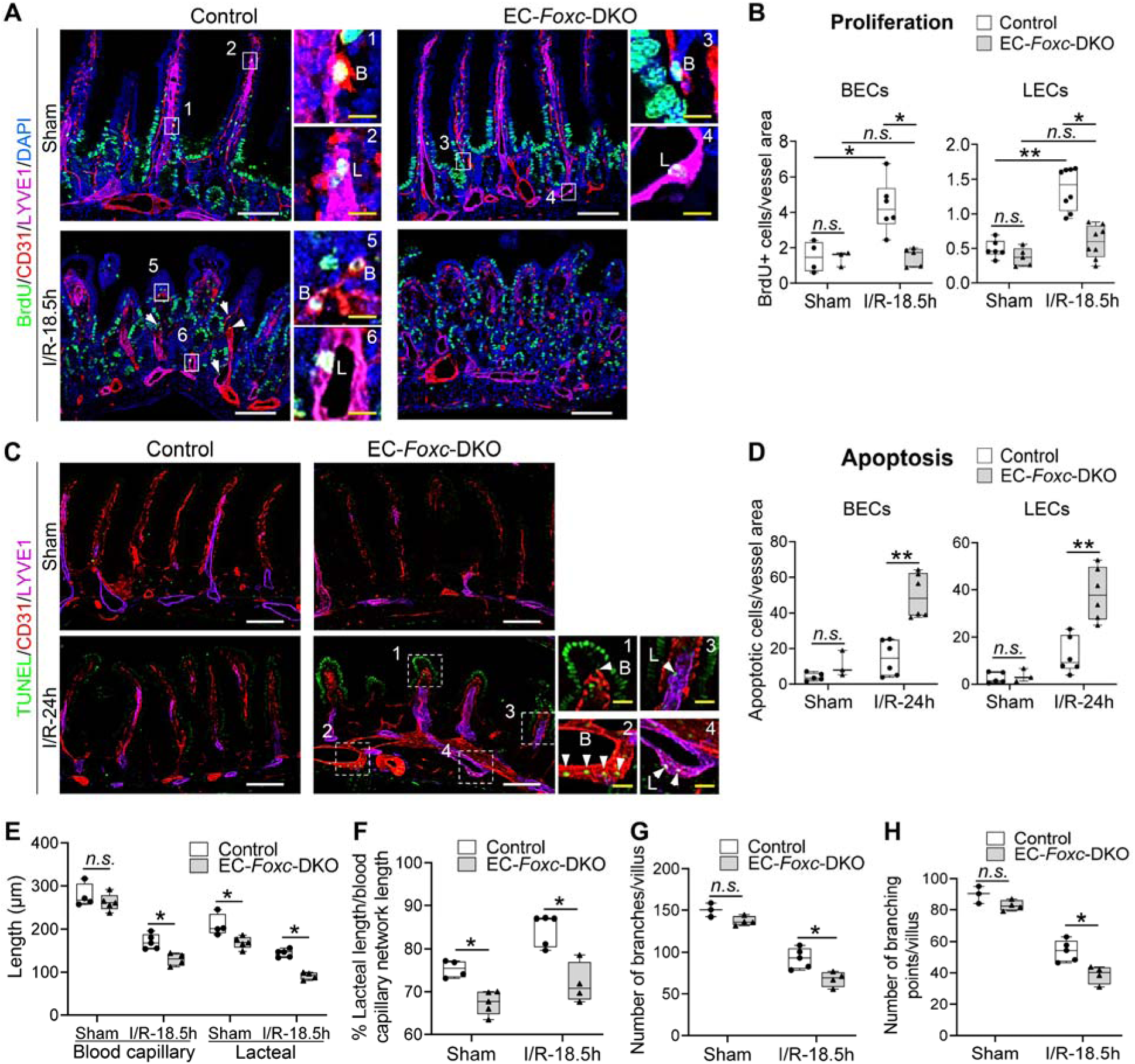
Impaired vascular regeneration after intestinal I/R injury in EC-*Foxc*-DKO mouse. (A) Representative BrdU/CD31/LYVE1/DAPI immunostaining images of intestinal paraffin sections (15 µm) from mice injected with BrdU 18.5h before euthanasia for the analysis of proliferative BECs (B in 1, 3, 5) and LECs (L in 2, 4, 6) in intestines. Arrow heads show BrdU+ BECs or LECs. White/yellow bars = 100 or 10 µm, respectively. **(B)** The numbers of BrdU+BECs and LECs per 0.1 mm^2^ blood vessel (CD31+LYVE1-) and lymphatic vessel (CD31+LYVE1+) area in intestinal mucosa were quantified respectively based on Figure 6A. Data are box-and-whisker plots, Kruskal-Wallis One-way ANOVA test, each symbol represents one mouse, N = 3∼8. **(C)** Representative TUNEL/CD31/LYVE1/DAPI immunostaining images of distal jejunums for the analysis of apoptotic BECs/LECs in intestinal paraffin sections (15 µm). High magnification images are from the dotted line boxes (1∼4). Arrow heads show the apoptotic BECs (B in 1, 2) and LECs (L in 3, 4) at villus (1, 3) or submucosa (2, 4). White/yellow scale bars = 100 or 20 μm, respectively. **(D)** The numbers of apoptotic BECs and LECs per 0.1 mm^2^ blood vessel (CD31+LYVE1-) and lymphatic vessel (CD31+LYVE1+) area in intestinal mucosa were quantified respectively based on Figure 6C. Data are box-and-whisker plots, Mann Whitney *U* test, each symbol represents one mouse, N = 3∼6. The length of blood capillary vasculature and lacteals were measured **(E)** based on Supplementary Figure 7, A and B. The percentage (%) of the lacteal length/blood capillary network length was then calculated **(F)**. The numbers of branches **(G)** and the branching points **(H)** of the villous blood vasculatures were counted based on the whole-mount staining of VEGFR2 (Figure 5A) as previously described ^66^ using ImageJ software. Data are box-and-whisker plots, Mann-Whitney U test, each symbol represents one mouse, N = 3∼5. \**P*<0.05, \*\**P*<0.01. *n.s.* = not significant.

Similarly, shorter lacteals were found in proximal jejunums in EC-*Foxc*-DKO neonatal mice at P7 after Tm treatment from P1 to P5 (Supplementary Figure 7, C-E), suggesting that *Foxc1/c2* are required for the maintenance of lacteal length. Furthermore, whole-mount immunostaining for VEGFR2 (Figure 5A) and subsequent quantification reveal a reduction in branches (Figure 6G) and branching points (Figure 6H) of blood capillaries in EC-*Foxc*-DKO mice after intestinal I/R injury. Together, these results suggest that EC-*Foxc1/c2* expression contributes to the repair of ischemic intestinal mucosa by promoting the recovery of BECs and LECs.

### *Reduced R-spondin 3* and *Cxcl12* expression in intestinal LECs and BECs of EC-*Foxc*-DKO mice, respectively, after I/R injury

To investigate molecular mechanisms associated with impaired intestinal regeneration in EC- *Foxc*-DKO mutants following I/R, we performed single-cell RNA sequencing (scRNA-seq) analyses of distal jejuna from control and EC-*Foxc*-DKO mice 18.5 hours after I/R injury. Dimensionality reduction and clustering analysis identified 22 transcriptionally distinct cell clusters (Figure 7A and Supplementary Figure 8A) based on known gene markers for each specific cell type (Supplementary Table 1), including BECs, LECs, stromal cells, epithelial cells and other cell types. When interpreted according to the recent classification of stromal cell populations in the ISC niche ^12^, the results from our scRNA-seq experiments indicates that *R-spondin 3* (*Rspo3*) is mainly expressed in two clusters (LECs and Telocytes/Trophocytes) after intestinal I/R injury (Figure 7B, cluster 12 and 20, respectively). Low expression of *Rspo3* was found in Myocytes/Pericytes cluster (Figure 7B, cluster 17). Further sub-clustering performed on Telocytes/Trophocytes cluster based on their known markers ^14^ identified 3 cell clusters, Trophocytes, Pdgfra^lo^ Cd81- stromal cells, and Telocytes (Supplementary Figure 8B). A dot plot used for visualizing differential gene expression (mean expression level) as well as gene expression frequency in different cell clusters (Figure 7C) showed that *Rspo3* was decreased in both LECs and Trophocyte clusters in EC-*Foxc*-DKO intestines compared with controls after I/R injury. In Trophocyte cluster, the moderately decreased trend of *Rspo3* was not significant (Supplementary Figure 8C). Similar *Rspo3* levels were found in the other 3 cell clusters between the two groups. Since the percentage of the LECs captured by scRNA-seq was very low (0.373%, Supplementary Figure 8A), the number of LECs obtained and used for analysis was very limited, and the decrease of *Rspo3* expression in LECs was not significant (Supplementary Figure 8D). However, by validating via IHC, RSPO3 protein was lower in intestinal LECs from EC-*Foxc*- DKO mice than control mice 24 hours after I/R injury (Figure 7D).

**Figure 7.**
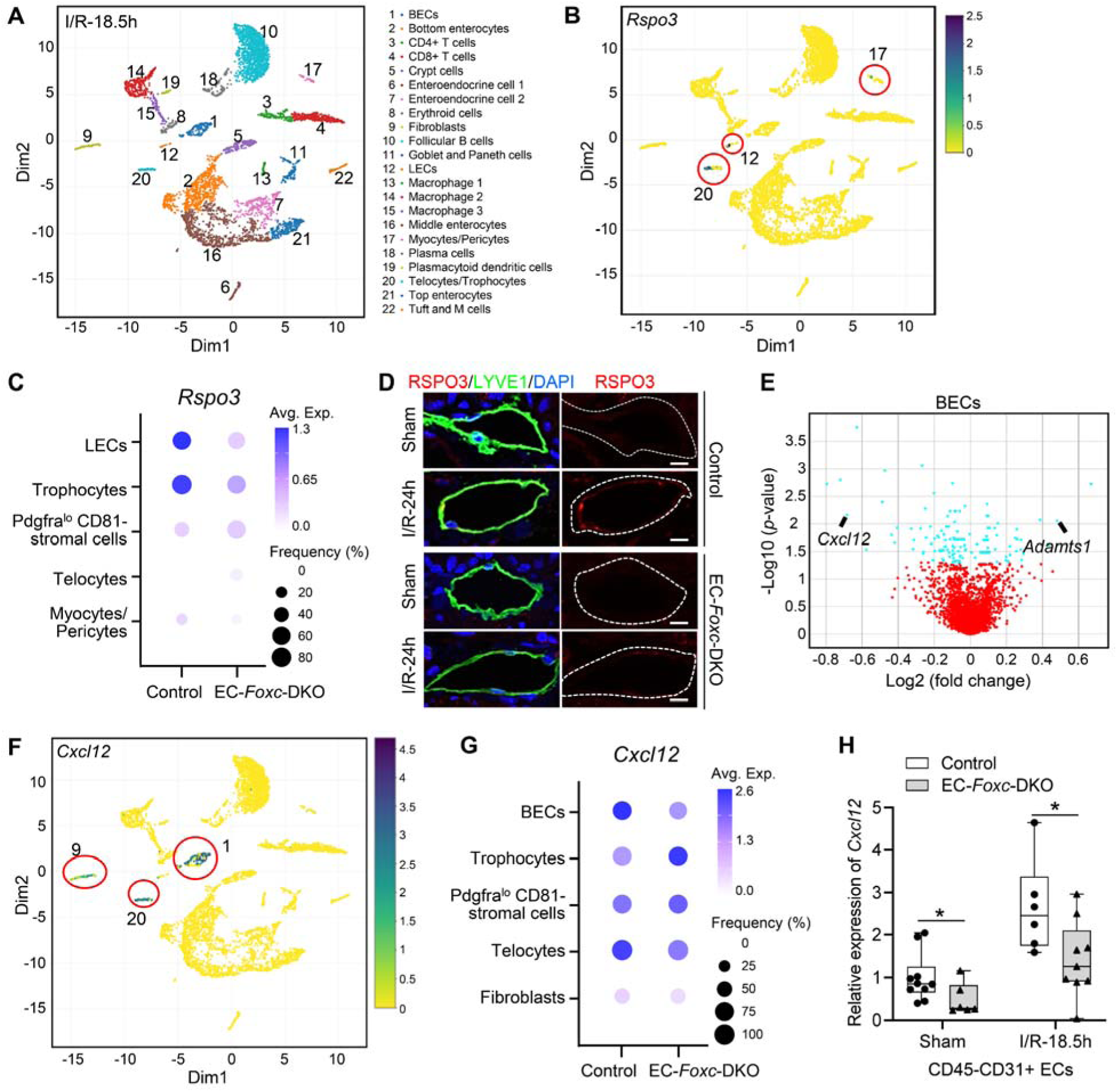
Single-cell RNA sequencing characterization of the distal jejunum from Control and EC-*Foxc*-DKO mice after I/R at 18.5h. (A) Visualization of unsupervised clustering of 22 distinct clusters by UMAP from the distal jejunum of both control and EC-*Foxc*-DKO mice after I/R at 18.5h. **(B)** UMAP visualization of *Rspo3* expression in LECs (12), Myocytes/Pericytes (17), and Telocytes/Trophocytes (20) cell clusters identified in Figure 7A. **(C)** Dot plot showing relative expression of *Rspo3* in 5 cell clusters identified by scRNA-seq. Fill colors represent normalized mean expression levels and circle sizes represent the within-cluster frequency of positive gene detection. **(D)** Representative images of lymphatic vessels (surrounded by the white dotted lines) in intestinal mucosa labeled with RSPO3/LYVE1. Scale bars = 10 µm. **(E)** Volcano plots of differential expression analysis of BECs in Control and EC-*Foxc*-DKO mice at I/R-18.5h using the MAST model. Blue dots denote genes with significant differential expression, *P* < 0.05. **(F)** UMAP visualization of *Cxcl12* expression in BECs (1), fibroblasts (9) and Telocytes/Trophocytes (20) cell clusters identified in Figure 7A **(G)** Dot plot showing relative expression of *Cxcl12* in 5 cell clusters identified by scRNA-seq. **(H)** Relative mRNA expression of *Cxcl12* in Dynabeads-isolated ECs (CD45-CD31+) from distal jejunum. Data are box-and-whisker plots, Mann-Whitney U test, each symbol represents one mouse, N = 6∼10, \**P*<0.05.

The results from our scRNA-seq experiments also identified numerous genes that were differentially expressed in intestinal BECs from EC-*Foxc*-DKO and control mice after I/R injury (Figure 7E), including the anti-angiogenic factor *Adamts1* ^47, 48^ and the CXC chemokine *Cxcl12*, which were up- and down-regulated, respectively, in EC-*Foxc*-DKO BECs. Notably, although *Cxcl12* was also expressed in both Telocyte/Trophocyte and Fibroblast clusters (Figure 7, F and G), it was significantly downregulated only in BECs (Figure 7, E and G), which have a larger cell population than the other two cell clusters (Figure 7F and Supplementary Figure 8A). Furthermore, qPCR analysis validated that *Cxcl12* expression was significantly reduced in isolated intestinal CD45^−^/CD31^+^ ECs of both sham- and I/R-treated EC-*Foxc*-DKO mice compared to their littermate controls (Figure 7H).

### R-spondin 3 treatment rescues the defective repair of small intestines in EC- and LEC-*Foxc*-DKO mice after I/R injury

RSPO3 is a key regulator of Wnt signaling during intestinal regeneration ^17, 18^, and RSPO3 prevents I/R-induced intestinal tissue damage and vascular leakage ^18^. Accordingly, we investigated whether RSPO3 treatment can rescue the defective repair of small intestines in EC- *Foxc*-DKO mice following I/R injury. Adult EC-*Foxc*-DKO mice were randomly assigned to two experimental groups and were treated with PBS or RSPO3 (5 μg in 100 μL PBS per mouse) by retro-orbital injection 30 min before intestinal ischemia. Quantification of Chiu scores was then performed 24 hours after I/R injury. RSPO3 treatment partially rescued the defective repair of small intestines in EC-*Foxc*-DKO mice (Figure 8, A and B). Given that RSPO3 is expressed in LECs of the small intestine, but not in BECs (Figure 7B), equivalent experiments were subsequently performed with adult mice carrying LEC-specific mutations of both *Foxc* genes. Similar rescue effects of the RSPO3 treatment were observed in LEC-*Foxc*-DKO intestines 24 hours after I/R injury with improvement of the Chiu score to the level similar to I/R-exposed control mice (i.e., grade 1) (Figure 8, C and D). Thus, treatment with RSPO3 is sufficient to fully recover intestinal I/R injury in LEC-specific *Foxc1*/*c2* deficient mice.

**Figure 8.**
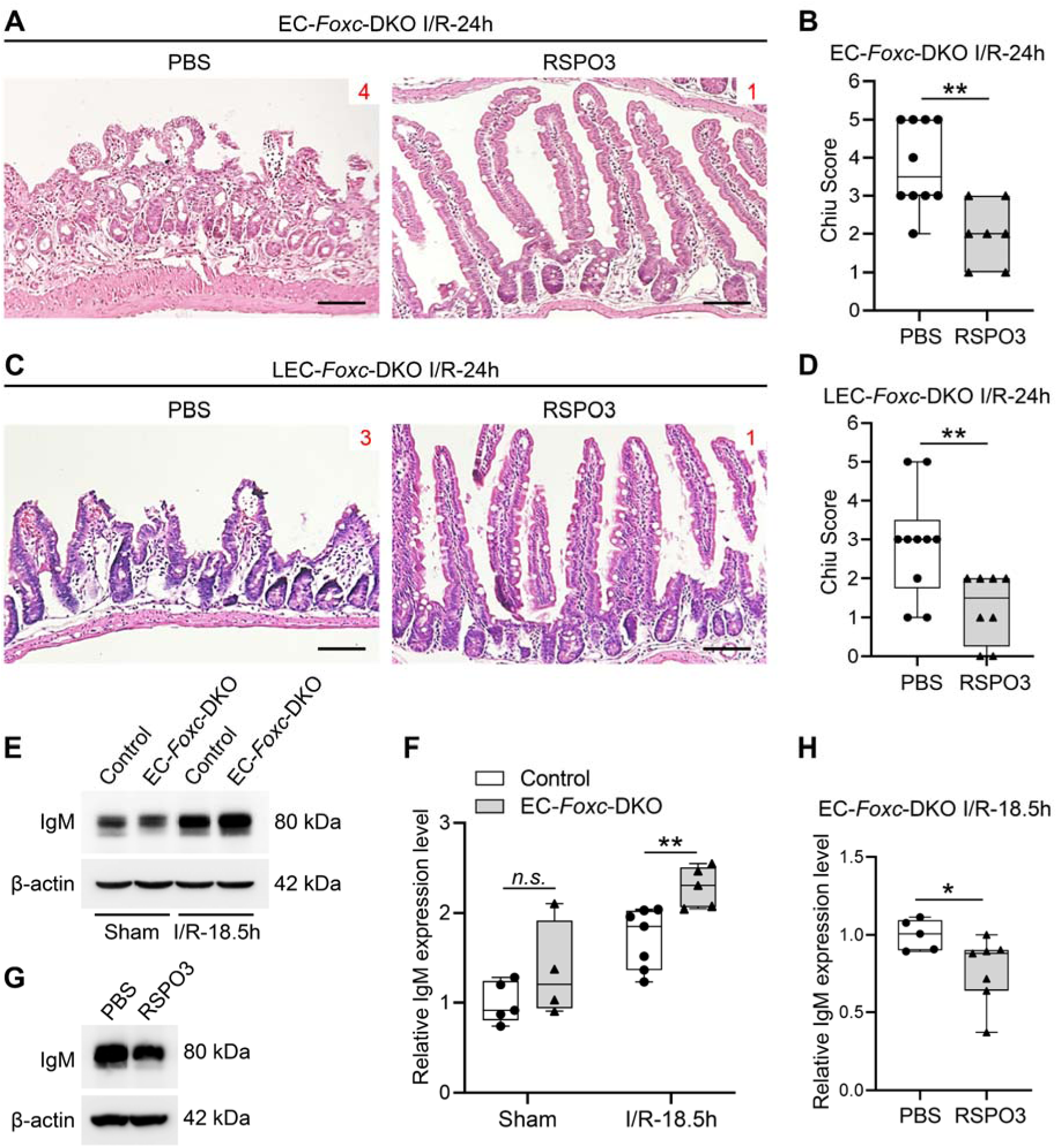
RSPO3 partially rescues impaired regeneration of intestinal mucosa in EC-*Foxc*- DKO and LEC-*Foxc*-DKO mice after I/R. In RSPO3 rescue experiment, each mouse was treated with 5 μg RSPO3 in 100μL PBS by retro-orbital injection 30 min before ischemia. PBS treated mice were used as control. Representative images of H&E staining show the rescue effects of RSPO3 in intestinal mucosa in EC-*Foxc*-DKO **(A)** and LEC-*Foxc*-DKO **(C)** mouse strains 24h after I/R. Red numbers indicate the Chiu scores. Scale bars = 100 μm. Quantification of Chiu Score for RSPO3 rescued **(B)** EC-*Foxc*-DKO and **(D)** LEC-*Foxc*-DKO intestines 24h after I/R. Data are box-and-whisker plots, Mann-Whitney U test, each symbol represents one mouse, N = 7∼13, \*\**P*<0.01. Representative Western blots (**E**, **G**) and densitometry measurements (**F**, **H**) show IgM (heavy chain) in intestinal tissue lysates from control and EC- *Foxc*-DKO mice in sham and after I/R at 18.5h (**E**, **F**) as well as IgM (heavy chain) in PBS- and RSPO3- treated EC-*Foxc*-DKO mice after I/R at 18.5h (**G**, **H**). Data are box-and-whisker plots, Mann-Whitney U test, each symbol represents one mouse, N = 4∼7, \**P*<0.05, \*\**P*<0.01. *n.s.*=not significant.

Immunoglobulin M (IgM) and complement also contribute to intestinal I/R injury ^18^, but RSPO3 treatment suppresses the deposition of IgM and complement in damaged intestinal tissue ^18^, likely by enhancing the activation of Wnt signaling in ISCs of the small intestine. Therefore, IgM deposition was evaluated in damaged intestines of control and EC-*Foxc*-DKO mice 24 hours after I/R injury via immunostaining (Supplementary Figure 9). Compared to control mice, IgM deposition was increased in EC-*Foxc*-DKO mice, whereas administration of RSPO3 reduced the level of IgM in EC-*Fox*-DKO mice. The levels of IgM deposition were further quantified via western blotting ^18, 31^. I/R-enhanced IgM levels in the intestinal tissues were greater in EC-*Foxc*-DKO mice compared to controls (Figure 8, E and F). More importantly, RSPO3 treatment significantly reduced the levels of IgM in the EC-*Foxc*-DKO mice after I/R injury (Figure 8, G and H). Collectively, these observations demonstrate that *Foxc1* and *Foxc2* expression in the intestinal vasculature contributes to intestinal mucosal recovery and regeneration after I/R injury.

### CXCL12 treatment partially rescue the defective repair of small intestines in EC-*Foxc*- DKO mice after I/R injury

*Cxcl12* was only downregulated in intestinal BECs, but not in the Telocyte/Trophocyte and Fibroblast clusters of EC-*Foxc*-DKO mutant intestine after I/R injury (Figure 7, E and G). Given evidence that hypoxia upregulates CXCL12 ^22, 23^ as a key regulator of angiogenesis ^24^, we examined whether pre-treatment with CXCL12 can rescue the defects in the vascular regrowth and intestinal repair associated with EC-*Foxc1/c2* deficiencies. Adult EC-*Foxc*-DKO mice were treated with PBS or CXCL12 via retro-orbital injection as described previously ^49^, and I/R induction was performed 30 minutes later. The degree of intestinal mucosal damage 24 hours after I/R injury was then quantified via the Chiu scoring system. CXCL12 treatment partially rescued the defective repair of small intestines in the EC-*Foxc*-DKO mice (Figure 9, A and B). Moreover, nuclear localization of β-catenin was observed in crypt ISCs of CXCL12-pretreated EC-*Foxc*-DKO mice after I/R injury compared to the PBS-treatment (Figure 9, C and D), and the number of cells expressing the Wnt target cyclin D1 (CCND1) ^13^ was significantly increased in the crypt ISCs of EC-*Foxc*-DKO mice treated with CXCL12 after I/R injury (Figure 9E).

**Figure 9.**
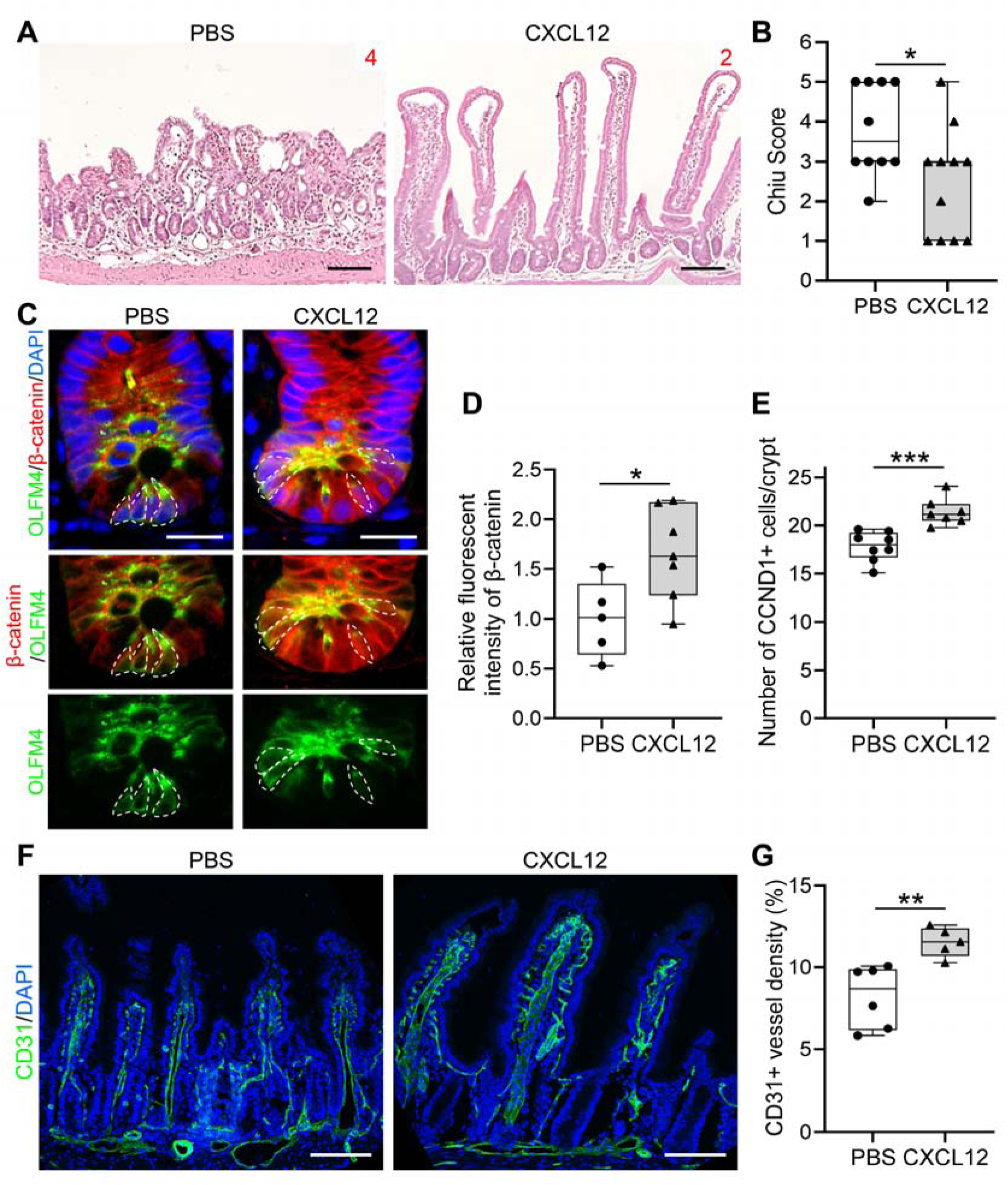
CXCL12 partially rescues impaired regeneration of intestinal mucosa in EC- *Foxc*-DKO mice after I/R. In CXCL12 rescue experiments, mice were treated with 50μg/kg CXCL12 in PBS by retro-orbital injection 30 min before ischemia. Mice treated with PBS were used as control. **(A)** Representative images of H&E staining show the rescue effects of CXCL12 in intestinal mucosa in EC-*Foxc*-DKO mice 24h after I/R. Red numbers indicate Chiu scores. Scale bars = 100 μm. **(B)** Quantification of Chiu Scores for CXCL12-rescued EC-*Foxc*-DKO mice 24h after I/R based on Figure 9A. Data are box-and-whisker plots, Mann-Whitney U test, each symbol represents one mouse, N = 10∼11, \**P*<0.05. **(C)** Representative images of crypts immunostained with OLFM4 and β-catenin in PBS- and CXCL12-treated EC-*Foxc*-DKO mice 24h after I/R. The total protein signal of β-catenin is up-regulated in CXCL12-treated crypts compared with PBS-treated group. The accumulation of β-catenin in the nuclei of ISCs (dotted circles) was found in CXCL12-treated mice but inhibited in PBS-treated mice. Paraffin sections (4 µm), scale bars = 20 μm. **(D)** Quantification of relative fluorescent intensity of β-catenin immunostaining within ISC based on Figure 9C. Data are box-and-whisker plots, Mann-Whitney U test, each symbol represents one mouse, N = 5∼7, \**P*<0.05. **(E)** Quantification of CCND1+ epithelial cells per crypt after I/R at 24h. Data are box-and-whisker plots, Mann-Whitney U test, each symbol represents one mouse, N = 7∼8. \*\*\**P*<0.001. **(F)** Representative confocal images of CD31 immunostaining of distal jejunums in PBS- and CXCL12- treated EC-*Foxc*-DKO mice after I/R at 24h. Paraffin sections (15 µm), scale bars = 100 μm. **(G)** Quantification of CD31+ vessel density (% = total CD31+ vessel area/total intestinal tissue area x 100%) based on Figure 9F. Data are box-and-whisker plots, Mann-Whitney U test, each symbol represents one mouse, N = 5∼6. \*\**P*<0.01.

Administration of CXCL12 also improved the defective vascular recovery (CD31+ vessel density) of EC-*Foxc*-DKO mice 24 hours after I/R injury (Figure 9, F and G). Therefore, CXCL12 treatment rescues the defects in vascular regrowth and intestinal repair associated with EC-*Foxc1/c2* deficiencies by enhancing vascular recovery and stimulating Wnt signaling in the crypts of the intestinal epithelium.

## Discussion

Little is known about the mechanisms that regulate expression of BEC/LEC-derived paracrine factors in tissue regeneration. In the present study, we demonstrate that EC-*Foxc1/Foxc2* expression is crucial for repair of the intestinal mucosa, BECs, and LECs after I/R injury, and that the EC- and LEC-*Foxc*-DKO mutations in mice impair canonical Wnt/β-catenin signaling in ISCs at the crypt base. Furthermore, our scRNA-seq data indicate that RSPO3 expression is attenuated in LECs and stromal cells of the EC-*Foxc*-DKO mice after intestinal I/R injury, which is at least partially attributable to impairments in intestinal regeneration because ISC activity appears to be crucially dependent on Wnt/β-catenin signaling in the subepithelial cellular microenvironment in the ISC niche ^10, 11^. We also show that CXCL12 is reduced in intestinal BECs of the EC-*Foxc*-DKO mice after I/R injury. Most significantly, treatment with RSPO3 and CXCL12 rescues the defective repair of small intestines in the LEC-*Foxc*-DKO and EC-*Foxc*- DKO mice, respectively. Consequently, this study elucidates the novel mechanisms that mediate the role of EC-specific FOXC1/C2 in repair of intestinal mucosa and vasculature, regulation of paracrine signaling factors, and activation of ISCs after intestinal I/R injury.

Adult single and compound EC- (or LEC)-specific *Foxc1/c2* mutant mice provide us with the *first* opportunity to comprehensively characterize how *Foxc* genes regulate the post-ischemic repair of intestinal blood/lymphatic vessels after I/R injury and intestinal epithelial regeneration by modulating Wnt/RSPO and CXCL12 signaling. Because the EC-*Foxc*-DKO and LEC-*Foxc*- DKO mutations are induced by different drivers (i.e., *Cdh5-Cre^ERT2^*and *Vegfr3-Cre^ERT2^*), the extent of *Foxc* downregulation in the two mutant mouse lines may not be equal. However, the degree of impairments in intestinal repair is consistently greater in the EC-*Foxc*-DKO mutant line than in the LEC-*Foxc*-DKO mutant line, suggesting that *Foxc1* and *Foxc2* are required in both BECs and LECs for intestinal tissue repair. Equivalent experiments were also performed with adult mice carrying EC- (and LEC-) specific KO mutations of each individual *Foxc* genes to determine the similarities and differences between the phenotypes associated with each deletion. While our qPCR and scRNA-seq analyses show that expression levels of *Foxc1* in intestinal ECs are higher than those of *Foxc2*, the phenotypic differences between EC- (or LEC-) *Foxc1*-KO and EC- (or LEC-) *Foxc2*-KO mice are not particularly distinct. Although the reason(s) for the phenotypic similarities remains unclear, recent evidence indicates that *Foxc2* is essential for the maintenance of intestinal LECs, and that treatment with antibiotics to deplete gut microbiota rescues the phenotype of LEC-specific *Foxc2* mutants, including lymphatic dilation in the small intestine ^50^. Bacterial translocation occurs when the intestinal barrier is disrupted by I/R injury, and I/R-induced intestinal damage can be attenuated by pretreatment of an antibiotic cocktail to deplete gut commensal bacteria ^31^. It remains to be elucidated whether commensal bacteria contribute to aberrant intestinal recovery and regeneration in our EC/LEC-specific *Foxc* mutant mice.

The EC-specific mutations of both *Foxc* genes in mice impair the regulation of RSPO3 in LECs (Figure 7), accompanied by reduced expression of ISC markers and Wnt target genes in crypt cells (Supplementary Figure 8E), and that RSPO3 treatment rescues the defective repair of small intestines in both EC-*Foxc*-DKO and LEC-*Foxc*-DKO mice after I/R injury (Figure 8). These observations are consistent with reports that RSPO3 prevents I/R-induced intestinal tissue damage ^18^, and that RSPO3 is a key regulator of Wnt signaling during intestinal regeneration ^17, 18^. While intestinal RSPO3 is known to be produced by LECs ^12, 19^, we demonstrate that pretreatment with RSPO3 almost completely rescues the impaired intestinal repair in the LEC- *Foxc*-DKO mice, implicating a paracrine effect of LEC-mediated RSPO3 signaling on the activation of ISCs after I/R injury. We analyzed data deposited in the ENCODE database ^51^ with HOMER software ^52^, as recently described in our analysis of the *PRICKLE1* locus ^41^, and identified putative FoxC binding sites (RYMAAYA or RYACACA) ^53,54,55^ in the vicinity of the human *RSPO3* locus (Supplementary Figure 10A and Methods). These sites are conserved between human and mouse, and contain histone methylated and acetylated regions (e.g., H3K4Me1 and H3K27Ac ChIP peaks), DNaseI hypersensitive regions, and transcriptionally active regions (identified by Transcription Factor ChIP-seq data). Thus, FOXC1 and FOXC2 are likely to regulate *RSPO3* expression directly in LECs. During our manuscript preparation, others have submitted bioRxiv preprints ^56,57,58^ showing that RSPO3 produced by intestinal LECs is required for maintaining ISCs in homeostasis and regeneration, reinforcing our finding that FOXC transcription factors mediate LEC-derived RSPO3 expression as a critical paracrine regulator in intestinal regeneration after I/R injury ^59^.

The chemokine CXCL12 is an angiocrine factor that is secreted from BECs and regulates organ- specific tissue repair, regeneration, and homeostasis, including the liver, bone marrow, and lung ^1, 60, 61^. However, the function of BEC-derived CXCL12 paracrine/autocrine signaling in damaged organs after injury remains largely unknown. The results from our study demonstrate that CXCL12 expression is downregulated in the intestinal BEC cluster and intestinal CD45-/CD31+ ECs of the EC-*Foxc*-DKO mice after I/R injury (Figure 7), and that CXCL12 treatment partially rescues the defective repair of small intestines in EC-*Foxc*-DKO mice after I/R injury (Figure 9). Our results are in accord with recent evidence that FOXC1 controls CXCL12 expression in CXCL12-abundant reticular (CAR) progenitors and Schwann cells ^62, 63^, and we also identified putative FOXC binding sites ^53,54,55^ in evolutionarily conserved regions of the human *CXCL12* locus (Supplementary Figure 10B and Methods). EC-derived CXCL12 promotes angiogenesis via an *autocrine* mechanism ^24^, and CXCL12-CXCR4 signaling cooperates with VEGF to enhance angiogenic processes such as the morphogenesis and sprouting of vascular tubes ^64^. Since VEGFR2 levels at the angiogenic front of growing blood capillaries in the villus are lower in EC-*Foxc*-DKO mice than in control mice after intestinal I/R injury (Figure 5A), it is likely that intestinal EC-derived CXCL12 autocrine signaling is defective in EC-*Foxc*-DKO mice. Consistently, impaired vascular recovery of the EC-*Foxc*-DKO mice after intestinal I/R injury is rescued by CXCL12 treatment (Figure 9). Since the vascular regrowth of villus BECs and LECs after I/R injury proceeds via a stepwise, interactive process, and regeneration of villus blood vessels precedes reconstruction of lacteals ^46^ (Figure 6 and Supplementary Videos), the severity of mucosal damage is greater in the EC-*Foxc*-DKO mice than in the LEC-*Foxc*-DKO mice. When the CXCR4 receptor in intestinal epithelial cells is activated by treatment with CXCL12- overexpressing MSCs, canonical Wnt/β-catenin is stimulated in the ISCs ^25^. Since CXCL12 treatment activates Wnt signaling in the ISC of the EC-*Foxc*-DKO mice (Figure 9), our results demonstrate that FOXC1 and FOXC2 promote endothelial-derived CXCL12 signaling in intestinal ischemia, thereby controlling vascular regrowth and intestinal regeneration.

In summary, our study demonstrates that BEC/LEC-FOXC1/FOXC2 expression regulates the repair of the intestinal vasculature and mucosal damage during intestinal regeneration after I/R injury by controlling the RSPO3 and CXCL12 signaling pathways. Thus, FOXC1 and FOXC2 regulate blood/lymphatic vascular function in the small intestine, as well as vascular mediated signaling to ISCs and the ISC niche during intestinal repair. Collectively, this study provides new insights into fundamental processes that are critically involved in recovery from ischemic disease and injury and may have important implications for the treatment of other ischemic conditions that are associated with impairments in tissue regeneration and stem-cell activity, including cardiovascular disease.

## Methods

### Animal husbandry

*Foxc1^fl/fl^*, *Foxc2^fl/fl^*, *Foxc1^fl/fl^;Foxc2^fl/fl^* ^39^, *Cdh5-Cre^ERT2^* ^40^, *Vegfr3-Cre^ERT2^* ^45^ and *Foxc2-Cre^ERT2^* ^33^ mice were used. EC-specific or LEC-specific *Foxc1*, *Foxc2*, and compound *Foxc1;Foxc2* mutant mice were generated by crossing *Foxc*-floxed females (*Foxc1^fl/fl^*, *Foxc2^fl/fl^*, and *Foxc1^fl/fl^;Foxc2^fl/fl^*) with *Cdh5-Cre^ERT2^;Foxc1^fl/fl^* (EC-*Foxc1*-KO), *Cdh5-Cre^ERT2^;Foxc2^fl/fl^* (EC- *Foxc2*-KO), *Cdh5-Cre^ERT2^;Foxc1^fl/fl^;Foxc2^fl/fl^*(EC-*Foxc*-DKO), *Vegfr3-Cre^ERT2^;Foxc1^fl/fl^* (LEC- *Foxc1*-KO), *Vegfr3-Cre^ERT2^;Foxc2^fl/fl^* (LEC-*Foxc2*-KO), *Vegfr3-Cre^ERT2^;Foxc1^fl/fl^;Foxc2^fl/fl^* (LEC-*Foxc*-DKO) males, respectively, as described previously ^41^. For Cre recombination efficiency detection, *mTmG/+;Cdh5-Cre^ERT2^;Foxc1^fl/fl^;Foxc2^fl/fl^*(mTmG/EC-*Foxc*-DKO), *mTmG/+;Vegfr3-Cre^ERT2^;Foxc1^fl/fl^;Foxc2^fl/fl^*(mTmG/LEC-*Foxc*-DKO) and *mTmG/+;Foxc2- Cre^ERT2^* mice were generated by crossing mTmG females (*mTmG/mTmG;Foxc1^fl/fl^;Foxc2^fl/fl^*and *mTmG/mTmG*) with EC-*Foxc*-DKO, LEC-*Foxc*-DKO and *Foxc2-Cre^ERT2^*males, respectively. Genotyping of mice was performed by Transnetyx Inc.

### Tamoxifen treatment

For adult mice, Tamoxifen (Tm, Cayman Chemical #13258) was dissolved in corn oil (Sigma #C8267) at 40 mg/mL by shaking at 37°C for 3-4 hours. 7–8-week-old male adult mice were treated with 150mg/kg Tm by oral gavage once daily for 5 consecutive days. For neonatal mice, each individual was treated with Tm (20 mg/mL, 75 μg) by oral gavage once daily from postnatal day 1 (P1) to day 5 (P5) ^41^.

### Cre recombination efficiency detection

mTmG/EC-*Foxc*-DKO, mTmG/LEC-*Foxc*-DKO and *mTmG/+;Foxc2-Cre^ERT2^* mice were used. The Cre negative mTmG mice were used as control. Twelve days after Tm treatment, the distal jejunum was harvested and fixed in 4% paraformaldehyde (PFA), followed by dehydration in 30% sucrose, and OCT (Sakura Finetek, USA) embedding. 10 or 15 μm cryosections were cut and immunostained with CD31 or LYVE1 antibody (Supplementary Table 2), counterstained with/without GFP antibody and a nuclear-specific dye DAPI. EGFP fluorescent signal was detected by imaging to evaluate the Cre recombination efficiency.

### Mouse small intestinal ischemia and reperfusion (I/R) surgery

Twelve days after Tm treatment, mice were subjected to small intestinal ischemia and reperfusion (I/R) surgery as previously described ^31^. Briefly, mice were anaesthetized with inhalation of Isoflurane. A midline laparotomy was performed, and the superior mesenteric artery (SMA) was then identified, isolated, and clamped by a small nontraumatic vascular clip. Heparinized-saline (500 μL, 10 U/ml) was added into the peritoneal cavity via syringe to avoid coagulation of blood. After this ischemic phase for 30 min, the clip was removed, and the intestine was allowed to reperfuse. 500 μL of sterile saline is administered to the peritoneal cavity to compensate for the fluid loss during surgery. A Chromic Gut Suture (4–0) was then used to close the muscle layer, followed by the skin closure with wound clips. Sham-operated mice were subjected to the exact same surgical procedure, aside from clip placement.

### BrdU treatment

Mice were treated with one dose of BrdU (Sigma #B5002, 10 mg/mL in PBS, 5 mg/kg) by intraperitoneal (i.p.) injection 2 h or 18.5h (for evaluation of proliferative intestinal epithelium or proliferative BECs and LECs, respectively) before tissue dissection (at I/R-24h and I/R-18.5h respectively).

### Tissue collection

Distal jejunum was selected for this study because this segment has the most severe mucosal damage after intestinal I/R surgery compared to other segments according to pilot experiments. Different time points (3 h, 4 h for an early injury stage, 18.5 h, 24 h and 48 h for a late repair stage) were chosen for different analysis purposes. For histological analysis, transcardial perfusion was performed on the adult mice with cold PBS followed by 4% PFA after anesthesia. Distal jejunum was harvested and cut longitudinally to expose the lumen. After several washes with PBS, the intestine was post-fixed in 4% PFA at 4°C for 4 h (for frozen or whole-mount samples) or for O/N (for paraffin embedded samples). For qPCR and Western blot on whole tissue lysates of the small intestine, blood was removed by transcardial perfusion with cold PBS. Distal jejunum was harvested, opened longitudinally, and washed with cold PBS, then snap-frozen in liquid nitrogen for the RNA isolation and protein extraction. For the tissue collection for neonatal intestinal whole-mount staining, the neonates were euthanized at P7 after Tm treatment from P1 to P5. The proximal jejuna were collected, washed with cold PBS, and fixed in 4% PFA at 4°C for 4 h, then subjected to the whole-mount staining protocol. Proximal jejunum was collected from neonatal mouse due to the ease of operation and similar lacteal length/blood capillary network length ratio between proximal and distal jejuna ^65^.

### Histopathological evaluation of intestinal mucosal damage

Paraffin sections of distal jejuna from the mice 24 h after intestinal I/R were stained with Hematoxylin and Eosin (H&E). Based on the H&E staining, Chiu Score ^42^ was used for evaluating the intestinal mucosal damage after I/R: grade 0, normal mucosa; grade 1, development of subepithelial Gruenhagen’s space at the apex of the villus; grade 2, extension of the space with moderate epithelial lifting from the lamina propria; grade 3, massive epithelial lifting with a few denuded villi; grade 4, denuded villi with exposed lamina propria and dilated capillaries; and grade 5, digestion and disintegration of the lamina propria, hemorrhage and ulceration. Higher scores represent more severe damage.

### Whole-mount (WM) staining

Whole-mount staining of small intestine was performed as previously described ^66^. Briefly, Distal jejunum was dissected, and fixed in fixative (0.5% PFA, 15% picric acid and 100 mM phosphate buffer, pH 7.0) at 4°C for O/N. Samples were washed with PBS, and subsequently dehydrated by 10% sucrose for 3 h, and 20% sucrose+10% glycerol in PBS for O/N at 4°C. After PBS-wash, samples were incubated with blocking buffer (5% donkey serum, 0.5% BSA, 0.3% Triton X-100, 0.1% NaN_3_ in PBS) for 2 h at 4°C, and then incubated with the indicated primary antibodies (Supplementary Table 2) diluted in the blocking buffer for O/N at 4°C. Samples were washed with PBST (0.3% Triton X-100 in PBS) for several times, followed by incubation with indicated fluorochrome-conjugated secondary antibodies (Supplementary Table 2) diluted in the blocking buffer for O/N at 4°C. The samples were washed again with PBST, post-fixed with 4% PFA, cut into thin strips (one or two villi wide), cleared with FocusClear (CelExplorer Labs #FC-101) and mounted on slides in mounting medium.

### Immunohistochemistry (IHC) staining

For IHC-P, 4 or 15 μm paraffin sections were deparaffinized, rehydrated, subjected to antigen retrieval, permeabilized with PBST, blocked with blocking buffer containing 5% Donkey serum in PBST for 30 min at room temperature (RT) and incubated with indicated antibodies (Supplementary Table 2) in blocking solution for O/N at 4°C. The sections were washed with PBS and incubated with indicated fluorochrome-conjugated secondary antibodies (Supplementary Table 2) in PBS for 1 h at RT. After wash with PBS, the sections were counterstained with DAPI and mounted with mounting medium. For IHC-F, 10 or 15 μm frozen sections were washed with PBS, and then subjected to the same blocking and antibody incubation protocols as IHC-P but without the antigen retrieval step. TUNEL staining was performed using *In Situ* Cell Death Detection Kit (Roche #11684795910) according to the manufacturer’s instructions.

### Imaging

H&E staining images were acquired using an Olympus Vanox AHBT3 Research Microscope (original magnification 100x, Tokyo, Japan). Fluorescent images were acquired using a Zeiss AxioVision fluorescence microscope or a Nikon A1 Confocal Laser Microscope with the software of Zeiss AxioVision SE64 Rel. 4.9.1 or NIS-Elements Viewer 4.20, respectively. Images were processed and analyzed with Adobe Photoshop, Imaris and Fiji (ImageJ) software. Imaris imaging software was used to create videos (Supplementary Videos, 1 and 2) for the 3-D structure of the blood and lymphatic vasculatures in villi after intestinal I/R.

### Rescue experiments

In rescue experiments, mice were treated with 5 µg RSPO3 (R&D systems #4120RS025CF) in 100 µL PBS ^18^, or CXCL12α (PeproTech #250-20A) in PBS at a dose of 50μg/kg of body weight (BW) ^49^ by retro-orbital injection 30 min before intestinal ischemia. Distal jejunum was harvested after I/R at 18.5 h (WB) or 24 h (H&E, IHC) for further analysis.

### Endothelial cell isolation from small intestine

ECs were isolated from the distal jejunum for further qPCR analysis. Briefly, the tissue was processed upon dissociating into single cell suspension in a digestion buffer (2 mg/mL Collagenase D, 0.2mg/mL DNAase I, 2 mg/mL Dispase II, 100 units/mL penicillin and 100 µg/mL streptomycin in HBSS) for 35 min at 37°C, followed by filtration through a 70μm and a 40μm cell strainer. After several washes, the cell suspension was then incubated with magnetic Dynabeads (Invitrogen #11035) labeled with CD45 antibody (Biolegend #103102) to deplete the CD45+ cell population. The CD45- cell suspension was then incubated with Dynabeads labeled with CD31 antibody (BD #553369) for the isolation of CD45-CD31+ cells. Finally, the CD45- CD31+ ECs were collected for RNA isolation and qPCR analysis.

### RNA isolation and qPCR analysis

A RNeasy Mini Kit (Qiagen #74104) was used for RNA extraction from isolated ECs. TRIzol^TM^ Reagent (Invitrogen #15596026) was used to isolate RNA from whole distal jejunum. The concentration of RNA was determined using NanoDrop™ 2000 Spectrophotometers (Thermo Scientific). cDNA was synthesized using an iScript reverse transcriptase kit (Bio-Rad #170- 8891). qPCR was performed on triplicates of cDNA samples by using QuantStudio® 3 Real-Time PCR System (Applied Biosystems), Fast SYBR reaction mix (Applied Biosystems), and gene specific primer sets. 18S was used as an internal standard for mRNA expression. Primer sequences are provided in Supplementary Table 3.

### Western Blot

The frozen intestinal tissue was grinned using mortar and pestle chilled with liquid nitrogen, followed by lysis in RIPA buffer (150mM NaCl, 1% Nonidet P-40, 0.5% sodium deoxycholate, 0.1% SDS and 50mM Tris, pH 7.4) containing protease inhibitors (Roche #4693116001). After centrifugation, the supernatant of the tissue lysates was collected and mixed with 5x Protein Loading Buffer. Equal amount of total protein for each sample was loaded and run on an SDS-PAGE gel. Samples were transferred to 0.45 µm nitrocellulose (Invitrogen) and Western blotted with the antibodies listed in Supplementary Table 2, followed by the reaction with ECL substrates. The chemiluminescent signal was then detected by imaging the blot with Azure c600 imaging system. Bands were quantified using ImageJ software.

### Preparation of single cell suspension from mouse small intestine

For single-cell RNA sequencing, mouse distal jejunums were collected 18.5h after intestinal I/R surgery. Two mice were used for each group: Control I/R-18.5h and EC-*Foxc*-DKO I/R-18.5h. Briefly, mice were anesthetized by isoflurane. Blood was removed by cardiac perfusion with cold PBS. The distal jejunum was then dissected, washed with cold PBS and cut into small pieces. The tissue was processed for scRNA-seq upon dissociating into single cell suspension in a digestion buffer as mentioned above for 35 min at 37°C, followed by filtration through a 70μm and a 40μm cell strainer. Cells were washed with washing buffer (0.5% BSA, 2mM EDTA in PBS, pH 7.4) for three times, and resuspended in PBS with 0.04% BSA at a concentration of 1200 cells/μL (according to 10x Genomics Document #CG00053 Rev B) before being passed through a 30 μm MACS SmartStrainer. The cell viability was tested by using the Cellometer Auto 2000 Cell Viability Counter (Nexcelom Bioscience, USA). The cell sample was processed for scRNA-seq only when the cell viability was more than 70%. Average cell viability for samples was determined to be 80.96%.

### Single cell 3’ gene expression library construction and sequencing

Single cell 3’ gene expression libraries were constructed by using the Chromium Next GEM Single Cell 3’ Reagent Kits v3.1 (10x Genomics, Pleasanton, CA, USA) according to the manufacturer’s manual CG000204 Rev D. The single cell libraries were assessed for quality (TapeStation 4200, Agilent, Santa Clara, CA, USA) and then run by using paired-end 50 bp sequencing on the Illumina HiSeq 4000 platform (Illumina, San Diego, CA, USA). 10,000 cells were targeted for each sample with a sequencing depth of 20,000 read pairs per cell.

### Pre-processing of single-cell RNA data

Following library generation and sequencing, raw sequencing data were de-multiplexed and mapped to the mouse reference genome (mm10) using the CellRanger toolkit (10X Genomics, version 2.1.0). Gene expression matrices were then generated from both Control and EC-*Foxc-* DKO mice using CellRanger. The matrix files were then utilized for data processing and downstream analysis using the BIOMEX browser-based software platform and its incorporated packages developed in R ^67^. Quality control and data pretreatment was performed in BIOMEX with the following manually set parameters: i) genes with a row average of <.001 were excluded for downstream analysis and ii) cells in which over 10% of unique molecular identifiers (UMIs) were derived from the mitochondrial genome were considered as dead cells and removed from downstream analysis. The data were then normalized in BIOMEX using similar methodology to the *NormalizeData* function as implemented in the *Seurat* package ^68^.

### Variable gene identification, dimensionality reduction, clustering analysis, and differential gene expression analysis

Following data pretreatment, BIOMEX was utilized for downstream dimensionality reduction of data and clustering analysis using the incorporated R packages. First, highly variable genes were identified utilizing the following feature selections: mean lower threshold = 0.01, mean higher threshold = 8, dispersion threshold = 0.5. Data (using highly variable genes only) was then auto-scaled and summarized by principal component analysis (PCA), followed by visualization using Uniform Manifold Approximation and Projection (UMAP; top 15 principal components (PCs)) to reduce the data into a two-dimensional space. Graph-based clustering was then performed in BIOMEX to cluster cells according to their respective gene expression profile using methodology similar to the *FindClusters* function in *Seurat* (clustering resolution = 0.5, k-nearest neighbors = 15). Marker set analysis was then performed in BIOMEX on highly variable genes to identify the top 10 gene markers expressed in each initial cluster using similar methodology described previously ^69^. Marker genes were then compared with single cell RNA-seq data from the small intestine of adult mice available from the Mouse Cell Atlas (MCA, http://bis.zju.edu.cn/MCA) and previously reported data ^70^ to identify transcriptionally unique cell populations. Clusters with highly similar expression patterns indicative of the same cell phenotype were merged into the same cluster. Differential gene expression analysis between Control and EC-*Foxc*-DKO mice for individual cell clusters was performed in BIOMEX using the Model-based Analysis of Single-cell Transcriptomics (MAST) package ^71^. *P*-values <.05 were considered statistically significant for differentially expressed genes.

### Data visualization

BIOMEX implementation of the *Plotly* software was used for UMAP and volcano plot visualization.

### Forkhead box C transcription factor binding prediction analysis in *RSPO3* and *CXCL12* loci

Putative FOX-binding sites in the *RSPO3* and *CXCL12* loci were first identified using the Hypergeometric Optimization of Motif EnRichment (HOMER) ^52^ suite of tools to scan the entire Genome Reference Consortium Human Build 37 (GRCh37 or hg19) genome corresponding to the conserved RYMAAYA FOX transcription factor binding motif or the reported RYACACA FOXC transcription factor binding motif ^72^. The output files were then uploaded to the UCSC genome browser ^51^ to identify putative binding sites corresponding to transcriptionally active areas denoted by histone modification, DNAse sensitivity, and additional transcription factor chromatin immunoprecipitation data as per work reported and summarized on the Encyclopedia of DNA Elements (ENCODE; https://genome.ucsc.edu/ENCODE/)(2012). Putative sites in the human genome were then searched against the Genome Reference Consortium Mouse Build 38 (mm10) genome using the Evolutionary Conserved Region (ECR) Browser (https://ecrbrowser.dcode.org) and rVista 2.0 tools to identify conserved and aligned putative binding sites between mouse and human sequences. Conserved and aligned putative FOX binding site sequences are underlined and bolded within the human *RSPO3* and *CXCL12* ECRs as shown in the Supplementary Methods.

### Mouse NEC model

24-hour-old mice were submitted to a well characterized NEC model ^35^ conducted in a 33-35°C infant incubator or left with the dam. The NEC protocol includes: 1) initial orogastric inoculation with a standardized adult mouse commensal bacteria preparation (10^8^ colony-forming units) and LPS (5 mg/kg) to perturb the normal intestinal colonization process; 2) gavage with formula every 3 h (Esbilac, 200 mL/kg/day); and 3) exposure to brief episodes of hypoxia (60 s in 100% N_2_) followed immediately by cold stress (10 min at 4°C) twice daily. This protocol induces intestinal injuries ranging from epithelial injury to transmural necrosis resembling human NEC which typically develop after 36 hours and has been widely used to study NEC pathogenesis ^73^. Pups were euthanized by decapitation at 24 hours into NEC and whole intestinal tissues were collected and fixed in formalin for 24 hours before tissue processing and paraffin block preparation.

### Quantification

For quantification of BrdU+ or CCND1+ cells in crypts, images from different fields of section under a 20x objective were acquired and about 15∼50 crypts were analyzed for each sample. The number of BrdU+ or CCND1+ cells was counted for each crypt. For quantification of β-catenin staining, about 5∼16 ISCs (OLFM4+) per sample were analyzed for the fluorescent intensity of β-catenin within the ISC by ImageJ. For analysis of BEC and LEC proliferation and apoptosis, confocal Z-stacks were acquired using a 20x objective from about 4∼ 8 different fields of 15 µm paraffin sections for each sample. Area of blood vessels (CD31+LYVE1-) and lymphatic vessels (CD31+LYVE1+) were measured using ImageJ software. Then the number of BrdU+ or TUNEL+ cells per 0.1mm^2^ vessel areas was calculated and compared between groups. Measurements for the length of blood capillaries and lacteals was performed as previously described ^66^. The numbers of branches and branching points for the villous blood vasculature was calculated based on the whole-mount staining of VEGFR2 as previously described ^66^. Around 30∼50 villi were analyzed for each sample. For CD31+ vessel density quantification, 5∼10 images were acquired using a 20x objective and the CD31+ vessel area and the intestinal tissue area in the images were measured using ImageJ software. CD31+ vessel density percentage was determined as the total CD31+ vessel area/total intestinal tissue area x 100%.

### Statistics

For quantification, statistical analysis was performed using GraphPad Prism 8.0 (GraphPad Software). *P* values were obtained by performing Mann-Whitney *U* test or Kruskal-Wallis One-way ANOVA test. Data are presented as box-and-whisker plots of representative experiments from at least three biological replicates. *P* values <0.05 were considered statistically significant. For scRNA-seq data, differential gene expression analysis between groups for individual cell clusters was performed in BIOMEX using the Model-based Analysis of Single-cell Transcriptomics (MAST) package ^71^. Adjusted *P* values <0.05 were considered statistically significant for differentially expressed genes.

### Study approval

All experimental protocols and procedures used in this study were approved by the Institutional Animal Care and Use Committee (IACUC) at Northwestern University.

## Supporting information

Supplementary Video 1_ Control IR-18.5h

Supplementary Video 2_EC-Foxc-DKO IR-18.5h

## Acknowledgments

We thank William Muller (Northwestern University) and Bona Jabri (University of Chicago) for helpful advice. *Cdh5-Cre^ERT2^*mice were kindly provided by Dr. Ralf Adams at the Max-Planck-Institute for Molecular Biomedicine, Germany. Single cell RNA-seq experiments were performed at the NUSeq Core Facility at Northwestern University.

## Author Contributions

CT, PRN, NU, XY, IGD, and TK designed and analyzed the experiments. CT, XY, TL, NU, and TL conducted the experiments. PRN analyzed scRNA-seq data. KA and SO provided Cre mouse lines. CT, PRN, and TK wrote the manuscript.

## Data and materials availability

All data needed to evaluate the conclusions in the paper are present in the paper and/or the Supplementary Materials. Additional data related to this paper may be requested from the authors.

