## Supplementary Materials for "Endothelial FOXC1 and FOXC2 promote intestinal regeneration after ischemia-reperfusion injury"

**This word file includes:**

Supplementary Methods
Supplementary Figure 1 to 10
Legends for Supplementary Video 1 and 2
Supplementary Table 1 to 3

**Other Supplementary Materials for this manuscript include the following:**

Supplementary Video 1 and 2

#### Supplementary Methods

**Human *RSPO3* and *CXCL12* Evolutionary Conserved Region (ECRs) with conserved and aligned putative FOX binding site sequences (underlined and bolded)**

*RSPO3 ECR-1.*>hg19 chr6:127441643-127442074

CTCCTGGAGCTCGCGAGGGGCAGGAAGCGATGCTGCTGCTCTGGGCTCTCGCACCTC

CGCGGCGTGCAGCCCAGGCCCAGCGGCCCCACCTGGTTTCCCAAGGGGATCCCTCTC

CCAGCGTCCCGCCCGGGAAGCCTTTCGGGGCAGACGTCGGGGCAGCGCGGAGGACA

CAACACCGGCTTGTGTGAGATGGAAATTTAGTAATCGCTTCTCGGCCGC**TGTTTGT**C

CCTGCCCTGCCCCTGGGCATGAGGGTCCCCTGAAAGCTCCCTTTGTATTGCTGGCCT

CGGCCAGACAAGGTTTGATTCACCTGAAAATTGCCTTTTAAAAGGTGGCAGTGCCA

GAGCGACAGGGGCCCTGGCCGGGCCTCACCGCCTTTAGGATCACCTAGCAGGGGGC

AGGAGAGCCCAAACTAAATATGTCAGGGCATCCGATTTATTATTCTTGGCCTCCGAA

TGGTGTGTACTCTGGTAACTCCCCACCCACTCTTTCATTCCGGCCCGGACGCGGAAG

GGGAGGCCTGGGGTTCAGCGGTGTCCACGCGCCTGGCGGTCCCCGCCTCCTCTCCTC

CCAGGGTCTGGCCTGGGGAATTGAGTCTCAGGGTCAGTGGGCATAGGGAGCCCGGG

CTGGAGCG

*RSPO3 ECR-2.*>hg19 chr6:127468065-127468506

TCTTGAAATGGATAATGATAGATTCCAAACACATGGAAATCTCTTGCCCCTTTTACTT

TTTAGGATCTTTGCAAGCTTACAATATGTACACGTTTTCTGTAAGTCACCAATGCTGA

GTTACTGGCATGAAAAATGACCCTGTTACTTGGAAAGTAGTTTCACTTACAAGTCCC

CCAGGCCCTGTAATGTCTAAACCTCCTGTGCCACTTTATGTGACTACCCCGCCCCCA

CAGAGGAGCATGCACAGGAAAAGCAGACTTCCCTTCCCCCACACATTTCCTTAGTT**T**

**<u>ATTTAC</u>**AAAACGTCTTGGAATGAGAATGAGCTGCTTGTGGTTCCTGTGGCTGATTCA
GGGATGGTTTCCTACAGGCAGAGGATGCTGGTCAACCGAATGACCTCTCTGTAACTA
ACCCGTGCACCCCTGTGGTAAGGCTGTTTGGTCTTATAGGTACCTCTTCTAACTAAGC
TTGGAGGGATTTGTTTTTGTGGTAAAGAACTTAGTAATAACCAAACGTCACTGTAAA
GACAGATTTAATAATGTTAAGGTCCATCAGAGCCTACTCCTTCTACTACCAACAAGA
GAAGCCAGAAATACACTGGGATGCCTTTAGATTCCTGTGCATCAATCTTTCTTTCTCT
AAGGATTATGG

*RSPO3 ECR-3*.>hg19 chr6:127497100-127497633

AATACCAGTAAATGGGGACACATTATATTGAATAAGGGTATTGTTAGCCAAATTCTA
AGATTCATCTTAAATTGTTTTCTTATAAGAATTGTGTATTTACCATTTTAAAAATCAC
TATTATTTTAAAACACTTAGAAAGTGAACATTTGAAAATGATGTGCCTTTGGATGCT
CTGTAATGTTAAGCAGATCCAGACATAAAGACAAAAGTAAATTCCAGAGTATTTTTG
TAGCCATGGAATCACCATAAAAAGGGGTTTTTGACCCCAATGTTACCGTAACATTGT
CTTCAGCATTTCATATTTAATTACAGTAGATTACTCACCAATATA**<u>ATAAATA</u>**TAGAT
TTATGAGATACTTTAATGTTCTAAAACAAATGAAAACCACCCAAGAGGAGCCTCACC
AAACCTGAGGTTGTCCAGATTGCATTGACTAAGATTAAGTAAAAGATCATTCATCTC
CAGAGGTCATGCAATTAATCTCAGAGTGGGAGTTAAAGCAATGACTAAGCAGAAAA
GGAAGCCAAATACAAGCTCGTAACAAAAGGTGCTGGGGCTCCAACATCAAGGAACT
TGTTATTTCCTTTTTATTTATTTATTTTTTTTTAATAGACCTAAAACACTCATTCCTTA
CTACTGGTTTCTTTGGGTCCTAAAATTCCACTTGGTTAGGTCAGCTATTTTCCATGAC
TATTTTTGATACGGTCAAACAAATACAAAGAATAAGCTTTTAAAAAAC

*RSPO3 ECR-4*.>hg19 chr6:127498181-127498446

TTCACCTTATTGAGATTCAAAACCTTGTGTTTTAAACCAAGAATAATTTTGAACTTGG

TCGTTCCTTATTGCAAGGACTTCCTCCAACTAGGGTGAATTTTCACCCACGATTAGCC

TGCCTTTGGAGTTTATGATGTGAAGTGCCACAGCTGATGGCCCATGAAAGAGACTGC

TAAGTCTATTGCTCACATCAATATTCCCGCTGGATATTTTGCTCATTCCTTGGGAATT

TCAAAGAATAACAAATA**ACCAACA**AATCATCAAAAACTGTAACATGTTACCCAATT

CTTGGGAATAAGCCAGTGAGACTAAAATCTGCTTAAGTATCACAAATGTGAATTCCC

AGCATTTCAAGCTTGTAAACTGTATCCAACACTCACCAATGTCTATTCCATTTCTTAG

CTGTACAGAAGGGGCACATTCATCTGGTAACTGTCAACAATGAGGTTCCTTCCTTCT

TGAACCC

*CXCL12 ECR-1*.>hg19 chr10:44888442-44889025

ATCAAAACCTTCACCTTTCTCTGCTGAAGGAATGGCCTTCTCTTATGGGCAGGGAGG

GTTTCCTAGGGAAAGCCCACCCAGGCAGGAGATGAGGAGAGCAGCATCTGAGCACA

CTTCATCCCACAGTGCCCATCCCATGAGTATCCTCCATAAATTACAAAGAAAGAAAA

AAAATAGGGAAAAAACAAACCTTTATTTCTC**TGTTTAC**TTCTCTGCATTTAATGAGC

AGTTGTTAACATGACTGAAACCATCTGATGATTTTTACCAAATGGAAAAATCTGCCT

ACAGGGGCAATAAA**ATAAATA**TTCAGAATAGAGAGAGGCAGTCATAAAAGACATT

ACCCGGTTGTAAACGGAGGCGGGTGGTGGTGATCTATTACCCCTGCCTCGGCAGCTT

TCAACAGAGTTCTGGAATTCCAGGAGGGGCCCTGACCCAAGGCAA**TATTTAC**TTCT

GCGGCTTCTTCATCAGGTCAGCATGGGTATAATTCTGTCTACCAGTTGACTGGAGCT

GAGGTTTCGAGCAGGAAGTGCAAACCCTGAGTGCTTATAACTCAGGAGGAGTGAGG

CACCCCTTCCCAGAGTATGCCAAGAAAAGCACATTGTACTGTCCTGGCTGCAGGGGT

GAGGCCCCTGCACACCCAGGCCATTATCAGCTTTGTGCCCTGGCCAACAGCGCCTCT

GGTTGGTGCATTTGTCAGCTGTATTTTACCCTAGAGCTCTGGGAGGCTCATCCTTTTT

TGGTATACCACCACGTGGAGAGAGCAGAGTTTTAATAGTGTGGCTGC

*CXCL12 ECR-2.*>hg19 chr10:44872457-44872993

GGGTTAAAAAAAAAAAAAAAAGATCCAAAAACTTGAGCTGCAGATCTAATCTGCTC

GTGAGAAAAGCCCATACACTGTCACACATGGGCTGTGAGAAGGGGTCTCAGACACC

TGACTGCAGGCAGGCTTAACTATATAAACCAGAAACGTCTATAAGCTCCATCACTAA

CAACTAATGAATTTTATTTCAG**GTAAATA**AATTCCCACATACAGTAGGACGTTTATA

CCATGAAACAATTAGCATTTTATTGCTAGTGCATATAATGTCACATTTGATACAATTT

TAGTACAAGTGAAAAAATACACTGTGGCTAACATTGAAAAGCTGCAATCACATTTAT

ATATCATATATATTTCTTTACAAATTGCCAGTAGTTTGAGATAATAGAGAAGTATAA

ACTACTGACATTCATATGGCTCCACTTCAAATATATGAATTGTTCGACT**ATAAATA**T

ATTTTGAAATACATTTGTTTTCTAAAGAAACGTAAAAAAAAATGTGCACAAAAATAT

ATATAAAAAAATGCCTTGCAAAAAGTTACAAATACCACCAGGACCTTCTGTGGATC

GCATTTATGCATGGAAATGTCACCTTGCCAACAGTTCTGATTGGAACCTGAAACCCT

GCTGTGGCTTCAGGAGGGGGTAGTGGCAAGATGATGGTTTATTCACTGATTTTTTCG

CTTCTGATTTCGGAAACCTCAGAGTTTGTTAGTGCCTC CATGGCATACATAGGCT

*CXCL12 ECR-3.*>hg19 chr10:44871130-44871571

GCGCAGGCCTTCTAAGAGAGGAAGTGGAGGGCGGGCTGGGGGGCTCAGCAACCTGG

GCATTCCTGGAGCTCCCAGGCTATTCTGGACAGAGTCCTGAGCACACAGGCTCCGCG

TCACAGACCCCCGCCCCAGTCTCTGCACAACTACTTTCTTCATAACTTAGCAGACCA

ACAAGGGCCTTAAAAAGCAC**ATAAATA**CAGAAGCCGGTGTGGCTGGCAGGGCAGC

AAAGCACTGCTCCCCCACGGAAGTCTGAGCCCCTGCCCAGTCTGCATGGGGGTCTGT

CCTGGAGGAGGATCGAGCAAATTTACAAAGCGCCGAGAGCAAGTGAACTGTGGTCC

ATCTCGAGGTGGCAGATAACTAGTTTTTCCTTTTCTGGGCAGCCTTTCTCTTCTTCTG

TCGCTTCTTTTTTCCTATCTTTTCTTTTTTCCCCACTTTTTCTTCTCTGCGCCCCCTTAG

ATAAAATTAGTAGAACCATTAATAATGTGGAAATAAACAAAAGTTCGTCTCAGTCTG

CATAAGATTTAACACTGGCCCGTGTACTGGTTAAACTGTGCCTTCAGTCTCAGGCCT

GGAGGCCCCTGGTCAAAGCACTGTTTATCAGTAATTCATTAAAATAAATTGGACATT

TCCTTCCATTCAAAC

*CXCL12 ECR-4.*>hg19 chr10:44868531-44868889

GCCCAGTCTCCCCACCTGCACAGCTCAGAGAATACAAAACCCAGGAGCCCTGAGTC

AGAGGAGTGGCTCCGTGGAAAGACACTCTAATAAGGCAAGTACAATAATGGCCTTA

GTCTAAGCTGCTACGTGTCGCCAGTGACACTGAATAATCAAACTAAATGAAAAATA

AATGTCATGTATTTTGCTACATAATCAAATGGTGAAAATATGGCAAAGTGTGCAAAA

CAAAGCCCTTGGCAATGCCTGGGGCCCTGCCCTGGCAGGGGAGGCTGTGGCAGGCC

CTTCCCTAACACTGGTTTCAGAGCTGGGCTCCTACTGTAAGGGTTCCTCAGGCGTCT

GACCCTCTCACATCTTGAACCTCC**TGTTGAC**AAGAACAGAGGGAAACACCATTAAG

GCAGGTTCCCCCCACACCCAGCCAGCTCCAGAAGGACAGCGGCCTGCGCGGAGGCC

CTGGGGCCAATCCCTACTTCCAAGTTGCCACCTTGGCCATGCATCTCCCTTCAGAGG

CCCTGCGACAACAGACGGGAGGGGAGTGCGAGA CGCAGCCAGGCTGAAGCGG

*CXCL12 ECR-5.*>hg19 chr10:44866547-44866769

GCAGCACCAGGTCCCGGAGGGACAGATGAGGGATCCCTAGTAAACAGCTCGTGGAC

GCACTTGACTAGCAGTTCGAAAGCAAAAAGAGATTCGGATTACAAGAGACTTTTCC

CTTGCAAATGGAAGACTGTATTTAAATGCATATTGCTTTAGCTGGGCAGTCTCCAGG

AAGGAGCTGATTGTTAGTAGAGGAATTGTTATGCAAATAATTTTCCCTGCAGTTTCT

ACTCTTATCTATTTTCTCCCATGGAGGAGAAATAGAAATGATTCCTTTCCTGACTGTA

GTCAAGATTTAGGGTCA**TGTTTGT**TTCCTCAACCAAGAATTCAATGGTCTTGGTTCA

TGGATTGGGGGCAGGTACATCCAAGTTCTACGTGACAGATTTTAAAATATCTTGGAT

TACCTGGATAATGACTGCCCCACTG

### Supplementary Figure 1

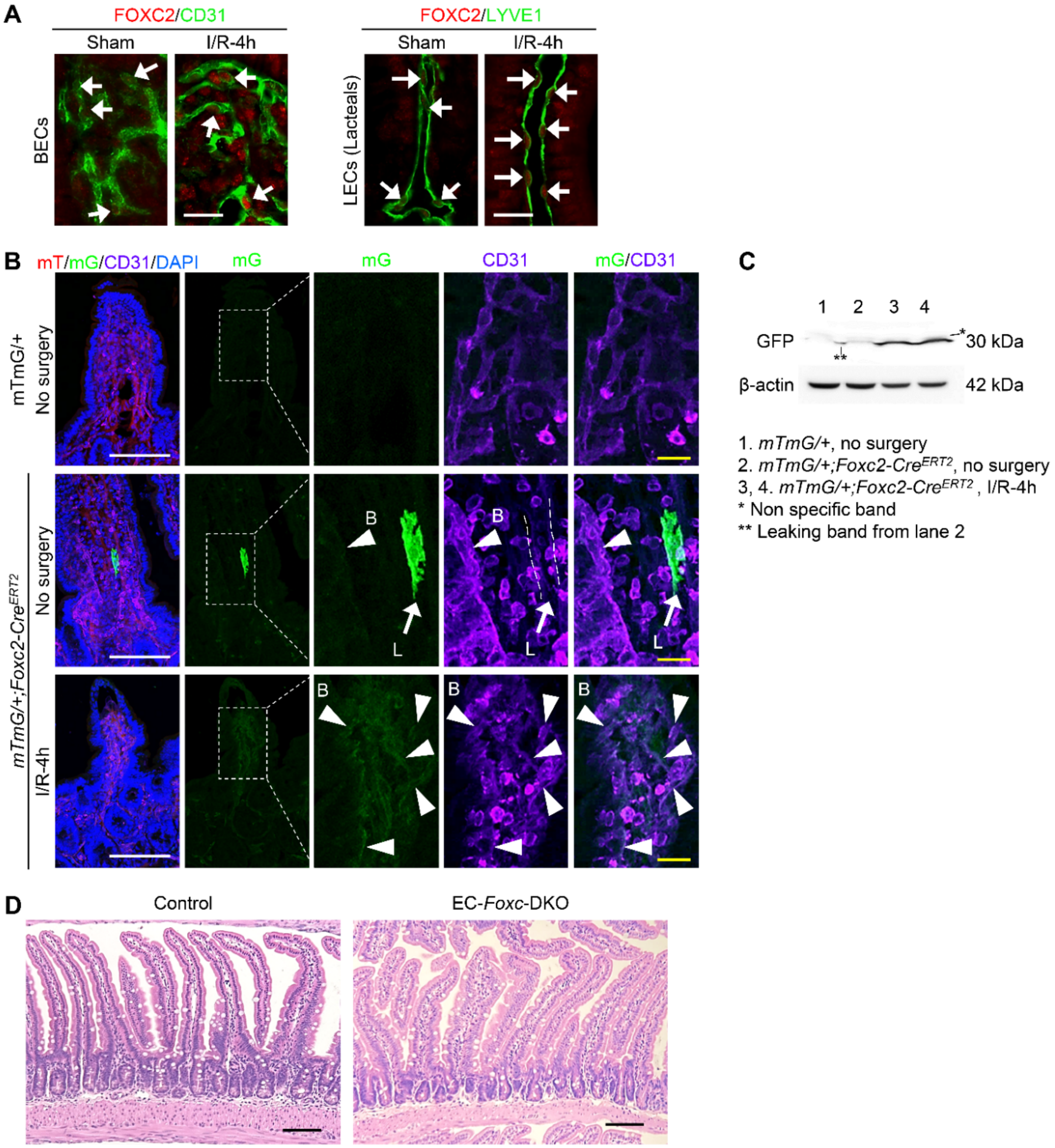


**Supplementary Figure 1. FOXC2 in intestinal ECs is up-regulated after I/R and down-regulated after Tm treatment in EC-*Foxc*-DKO mice. (A)** Representative immunostaining images of villi show FOXC2 is up-regulated in intestinal BECs (CD31+) and LECs (LYVE1+) after I/R at 4h in control adult mice (*Foxc1$^{f/f}$;Foxc2$^{f/f}$*). **(B)** Representative intestinal mucosal images of CD31 immunostaining with mT/mG signals on frozen sections (15 μm) in *mTmG/+;Foxc2-Cre$^{ERT2}$* mice without surgery or 4h after I/R. *mTmG/+* mice without surgery were used as control. Lacteals (L, outlined by dashed lines) express FOXC2-GFP (arrows) after Tm treatment. FOXC2-GFP is up-regulated and weakly expressed in villous blood vasculatures (B with arrow heads) after I/R at 4h. White scale bars = 100 μm. Yellow scale bars = 20 μm. **(C)** Representative Western blots show the up-regulation of FOXC2-GFP in intestinal lysates of *mTmG/+;Foxc2-Cre*$^{ERT2}$ mice 4h after I/R compared with the mice without surgery. **(D)** Representative H&E staining images of intestinal mucosa in Control and EC-*Foxc*-DKO mice after Tm treatment without surgery. Scale bars = 100 μm.

#### Supplementary Figure 2

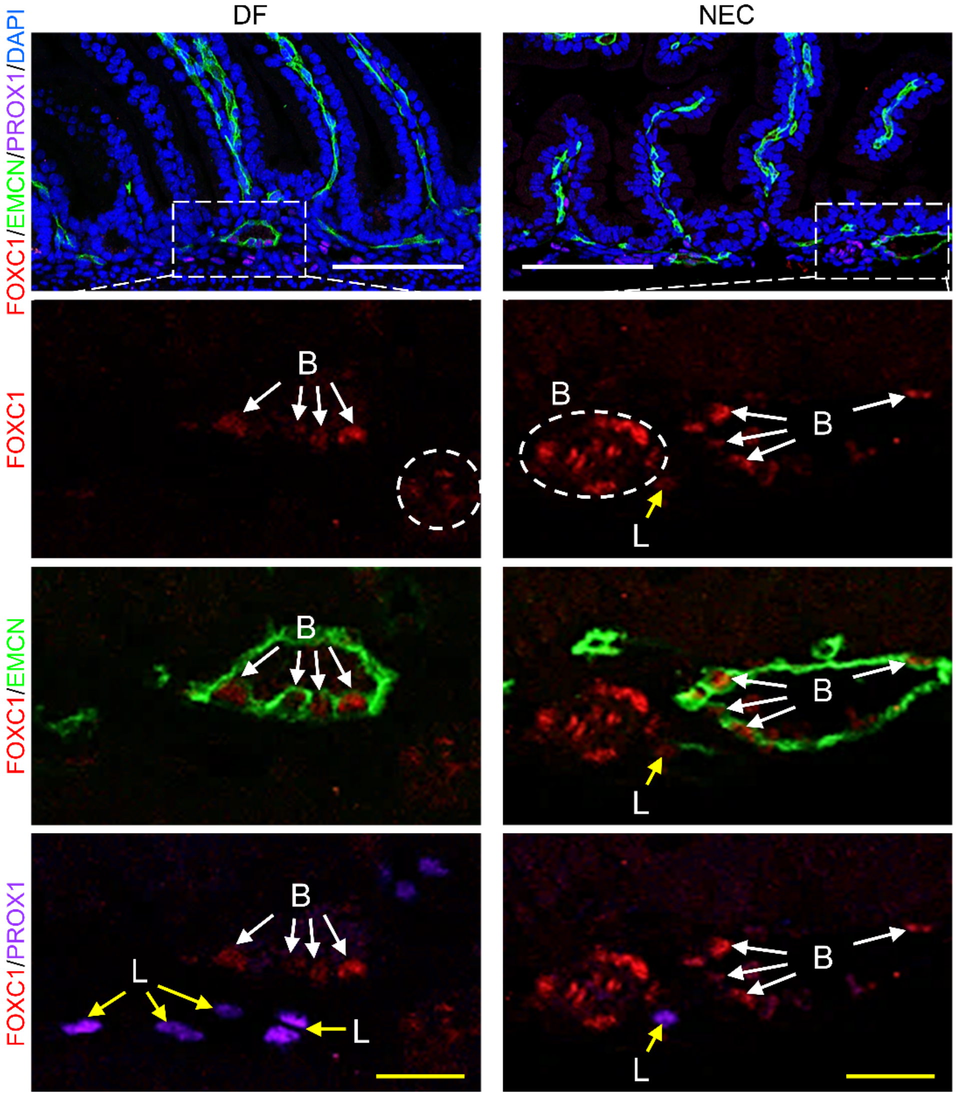


**Supplementary Figure 2. FOXC1 is increased in BECs and LECs in mouse NEC model.** Immunostaining of FOXC1/EMCN/PROX1/DAPI was performed on paraffin sections (4 µm) of small intestines from 2-day old neonatal mice 24 hours after being subjected to the necrotizing enterocolitis (NEC) protocol [35] . Dam-fed (DF) pup littermates were used as control. The level of FOXC1 is increased obviously in BECs (EMCN+, B with white arrow; as well as circled area) in NEC intestine compared with DF intestine. FOXC1+ cells in circled area are arterial BECs with EMCN-. FOXC1 can be found weakly expressed in the LECs (L with yellow arrow, PROX1+) in NEC intestine but is hardly detectable in LECs in DF intestine. PROX1 is a nuclear marker for LECs. White/yellow scale bars = 100 µm or 20 µm, respectively.

#### Supplementary Figure 3

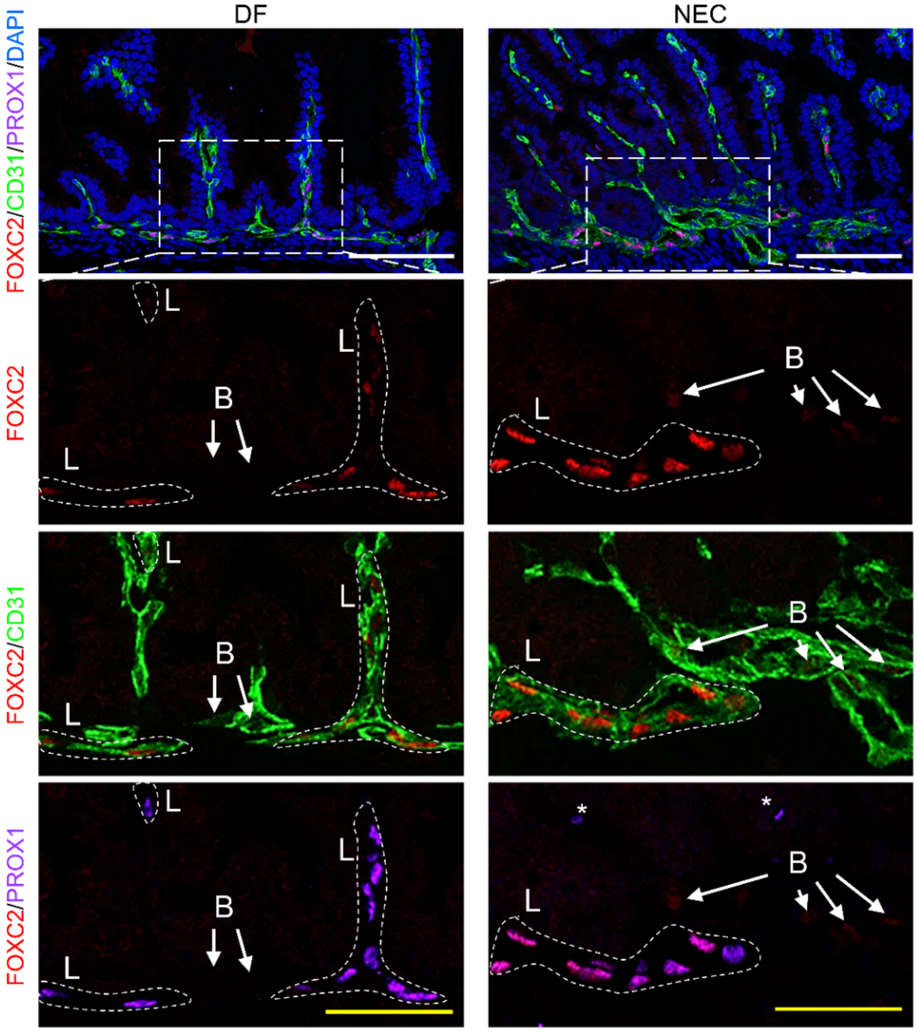


**Supplementary Figure 3. FOXC2 is increased in LECs and BECs in mouse NEC model.** Immunostaining of FOXC2/CD31/PROX1/DAPI was performed on paraffin sections (4 μm) of small intestines from 2-day old neonatal mice 24 hours after being subjected to the necrotizing enterocolitis (NEC) protocol [35]. Dam-fed (DF) pup littermates were used as control. FOXC2 can be detected in LECs (L, CD31+PROX1+; circled) in DF intestine, and is increased obviously in LECs in NEC intestine compared with DF intestine. FOXC2 can be found weakly expressed in the BECs (B, CD31+PROX1-) in NEC intestine (white arrow) but is hardly detectable in BECs in DF intestine (yellow arrow). CD31 is a marker for both BECs and LECs. PROX1 is a nuclear marker for LECs. *: non-specific staining of PROX1. White/yellow scale bars = 100 μm or 50 μm, respectively.

#### Supplementary Figure 4

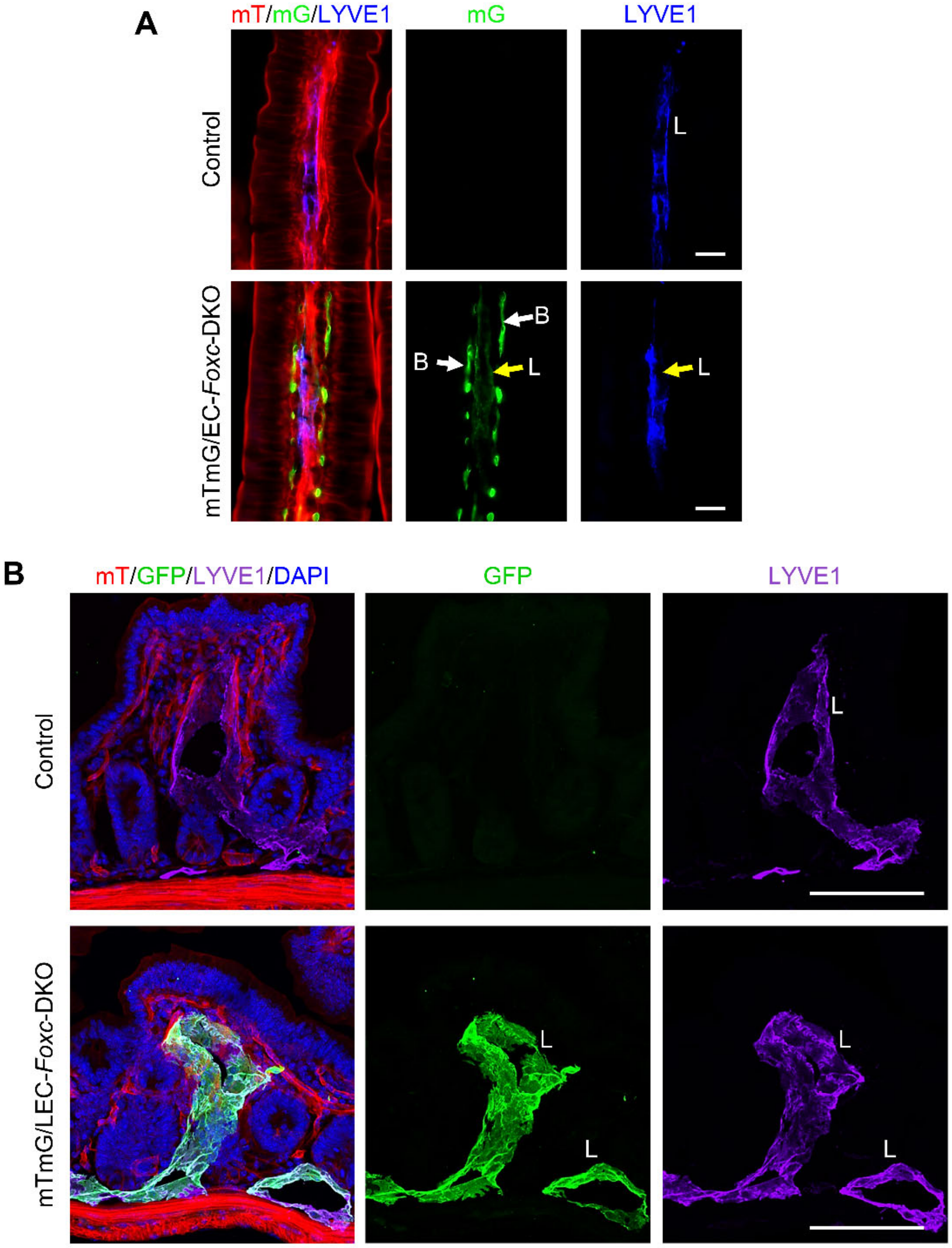


**Supplementary Figure 4. Cre recombination efficiency detection in EC-*Foxc*-DKO and LEC-*Foxc*-DKO mouse strains. (A)** Mice (Control: *mTmG/+;Foxc1$^{f/f}$;Foxc2$^{f/f}$*, mTmG/EC-*Foxc*-DKO: *mTmG/+;Cdh5-Cre$^{ERT2}$;Foxc1$^{f/f}$;Foxc2$^{f/f}$*) were treated with 150 mg/kg Tm by oral gavage for 5 days. Seven days after Tm treatment, the distal jejunum was collected and the frozen sections (10 µm) were stained with LYVE1 for the detection of GFP signal in blood vessels (B, LYVE1-GFP+) and lacteals (L, LYVE1+GFP+). Scale bars = 20 µm. **(B)** Mice (Control: *mTmG/+;Foxc1$^{f/f}$;Foxc2$^{f/f}$*, mTmG/LEC-Foxc-DKO: *mTmG/+;Vegfr3-Cre$^{ERT2}$;Foxc1$^{f/f}$;Foxc2$^{f/f}$*)

were treated with 150 mg/kg Tamoxifen by oral gavage for 5 days. Twelve days after Tm dose, the distal jejunum was collected and the frozen sections (15 μm) were stained with GFP and LYVE1 antibodies. Confocal images show the VEGFR3-GFP is expressed in the LYVE1+ lymphatic vessels (L). Scale bars = 100 μm.

#### Supplementary Figure 5

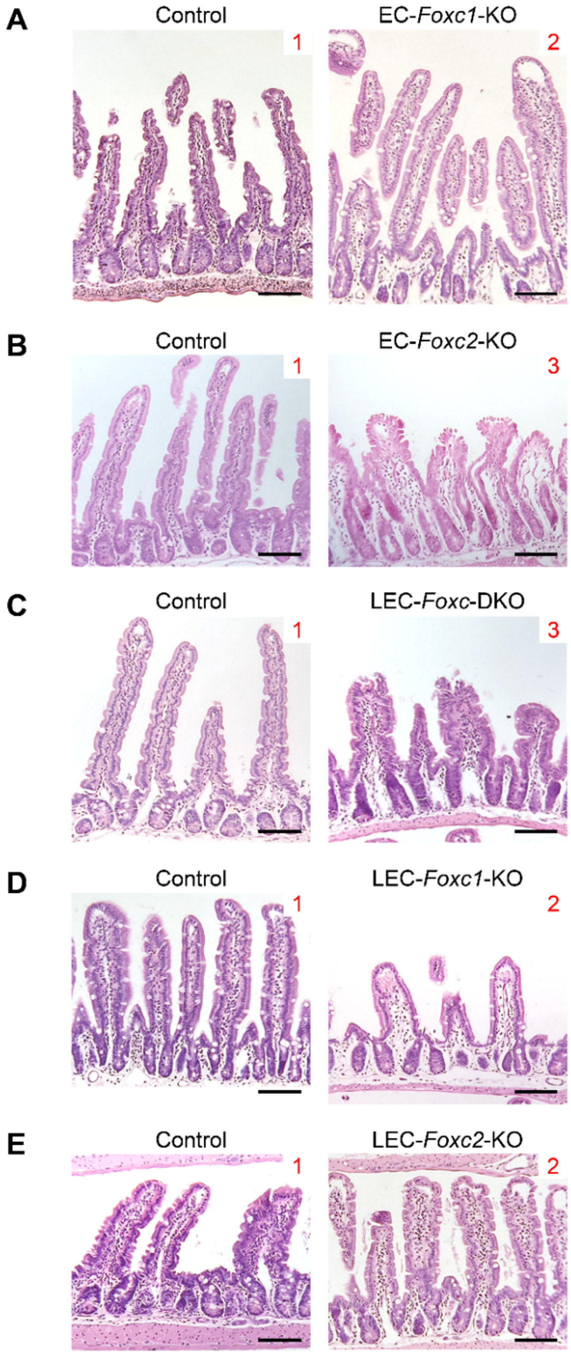


**Supplementary Figure 5. H&E staining for different mouse strains.** Representative H&E staining images of the distal jejuna 24 h after I/R in different mouse strains: **(A)** EC-*Foxc1*-KO, **(B)** EC-*Foxc2*-KO, **(C)** LEC-*Foxc*-DKO, **(D)** LEC-*Foxc1*-KO, **(E)** LEC-*Foxc2*-KO and their control mice. The intestinal ischemic injury grading in the Chiu scoring system is indicated by red numbers (0~5). Scale bars = 100 μm. The quantification of Chiu Score for these mouse strains are shown in Figure 2, D and E, Figure 4, A-C, respectively.

#### Supplementary Figure 6

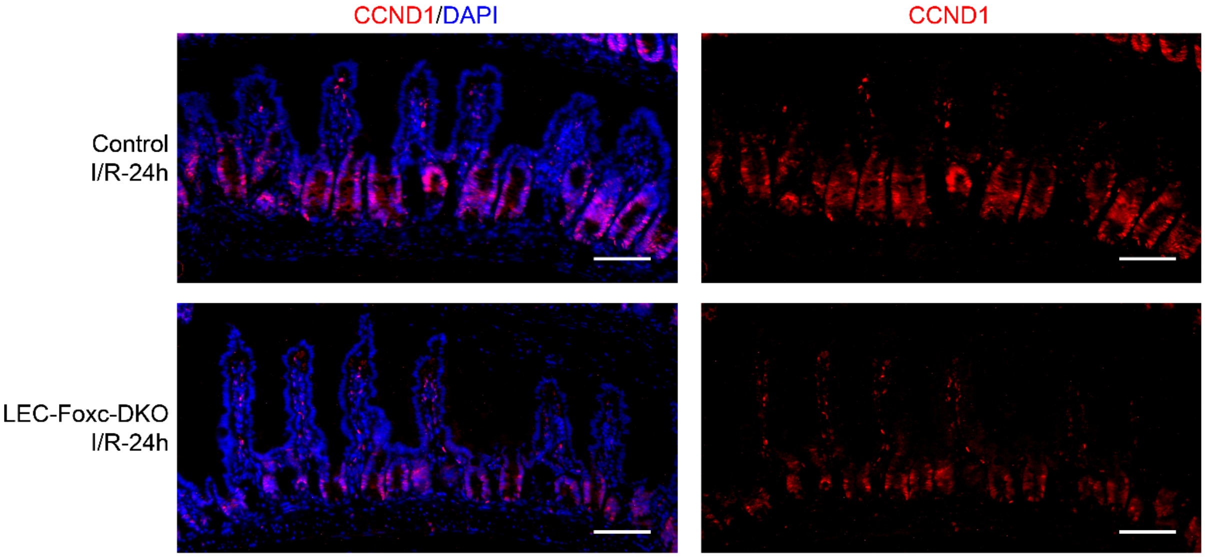


**Supplementary Figure 6. Immunostaining of CCND1 in intestinal mucosa in LEC-*Foxc*-DKO mice.** Representative images of intestinal mucosa labeled with Cyclin D1 (CCND1) in LEC-*Foxc*-DKO mice compared with Control group 24h after I/R. Scale bars = 100 µm. Quantification data for CCND1+ epithelial cells per crypt are shown in Figure 4G.

#### Supplementary Figure 7

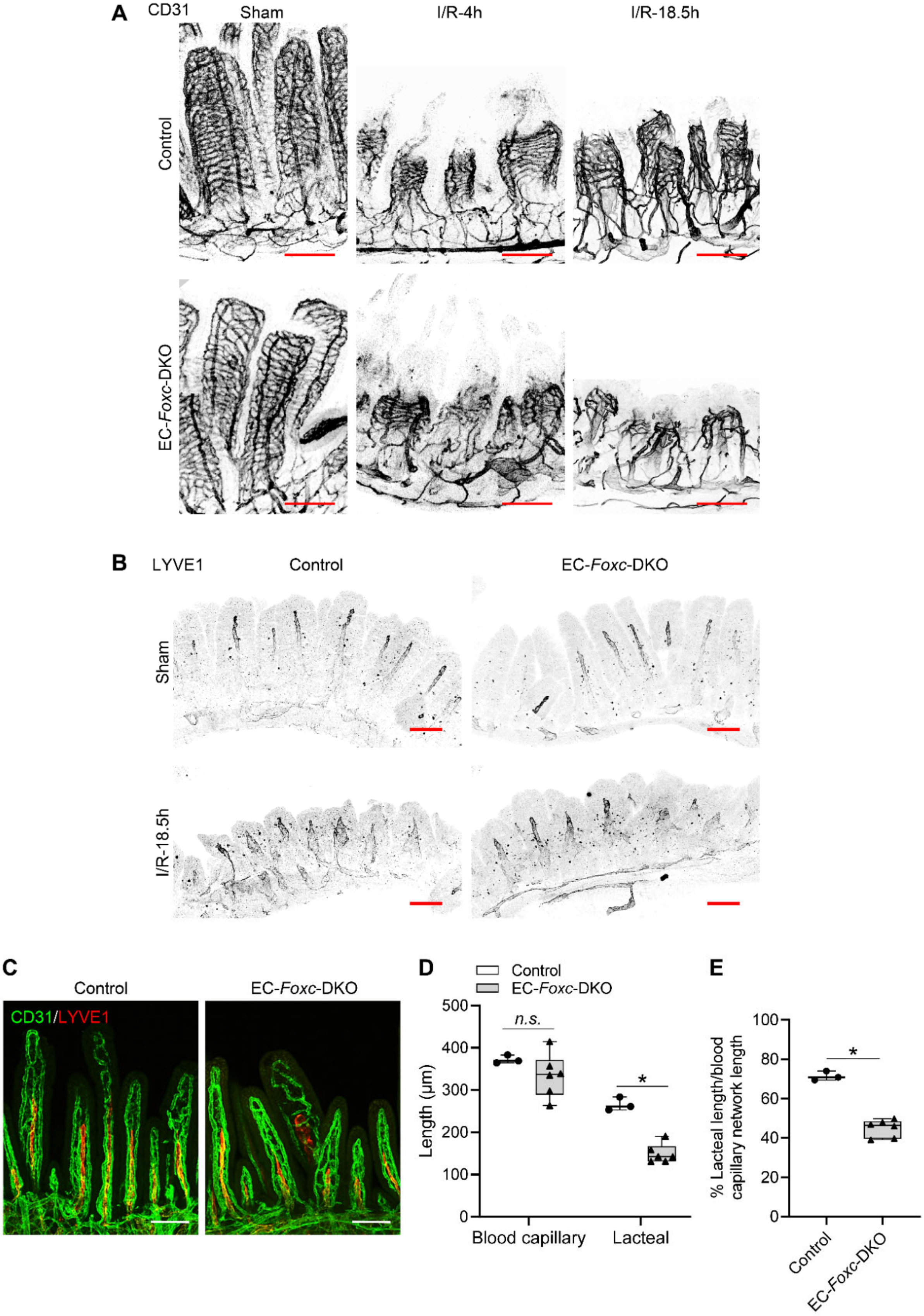

**Supplementary Figure 7. EC-iKO of *Foxc1/c2* results in impaired repair of blood and lymphatic vasculatures in intestinal villi after I/R.** Representative images of whole-mount distal jejuna stained with CD31 **(A)** and LYVE1 **(B)** show the damage of blood **(A)** and lymphatic vasculatures **(B)** in control and EC-*Foxc*-DKO villi at 4h and/or 18.5h after reperfusion. Scale bars = 100 µm. **(C)** Representative whole-mount proximal jejuna immunostained with CD31 (green) and LYVE1 (red) in neonatal mice treated with Tm from P1 to P5 and euthanized at P7. The length of blood capillary vasculature and lacteals were measured **(D)** based on Supplementary Figure 7C. The percentage (%) of the lacteal length/blood capillary network length was then calculated **(E)**. Data are box-and-whisker plots, Mann-Whitney U test, each symbol represents one mouse, N = 3~6. *$P$<0.05, *n.s.* = not significant.

**Supplementary Figure 8**

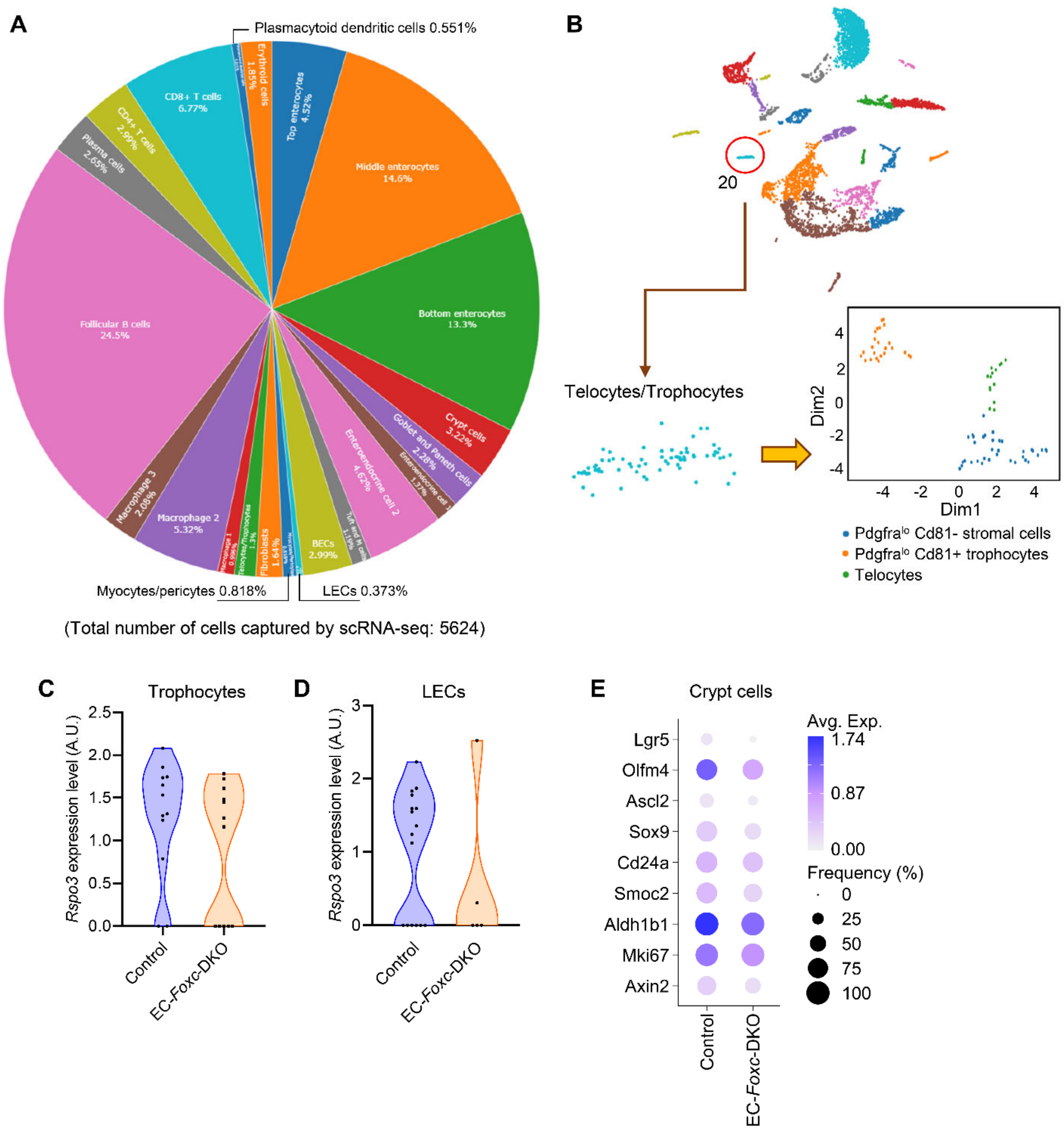


**Supplementary Figure 8. ScRNA-seq analysis to the small intestines from control and EC-*Foxc*-DKO mice 18.5h after I/R. (A)** Pie chart showing the percentage of each cell clusters identified in Fig. 7A of total cell population. **(B)** Sub-clustering performed on the cluster 20 (Telocytes/Trophocytes). **(C-D)** Violin plots of the *Rspo3* expression in trophocytes **(C)** and LECs **(D)** in intestine at I/R-18.5h. **(E)** Dot plot showing relative expression of different genes identified by scRNA-seq were decreased in crypt cell cluster in EC-*Foxc*-DKO mice compared with control mice at I/R-18.5h. Fill colors represent normalized mean expression levels and circle sizes represent the within-cluster frequency of positive gene detection. *Lgr5*, *Olfm4*, *Ascl2*, *Sox9*, *Cd24a*, *Smoc2* and *Aldh1b1* are ISC markers. *Mik67* is proliferative marker. *Ascl2*, *Sox9* and *Axin2* are Wnt/β-catenin target genes.

#### Supplementary Figure 9

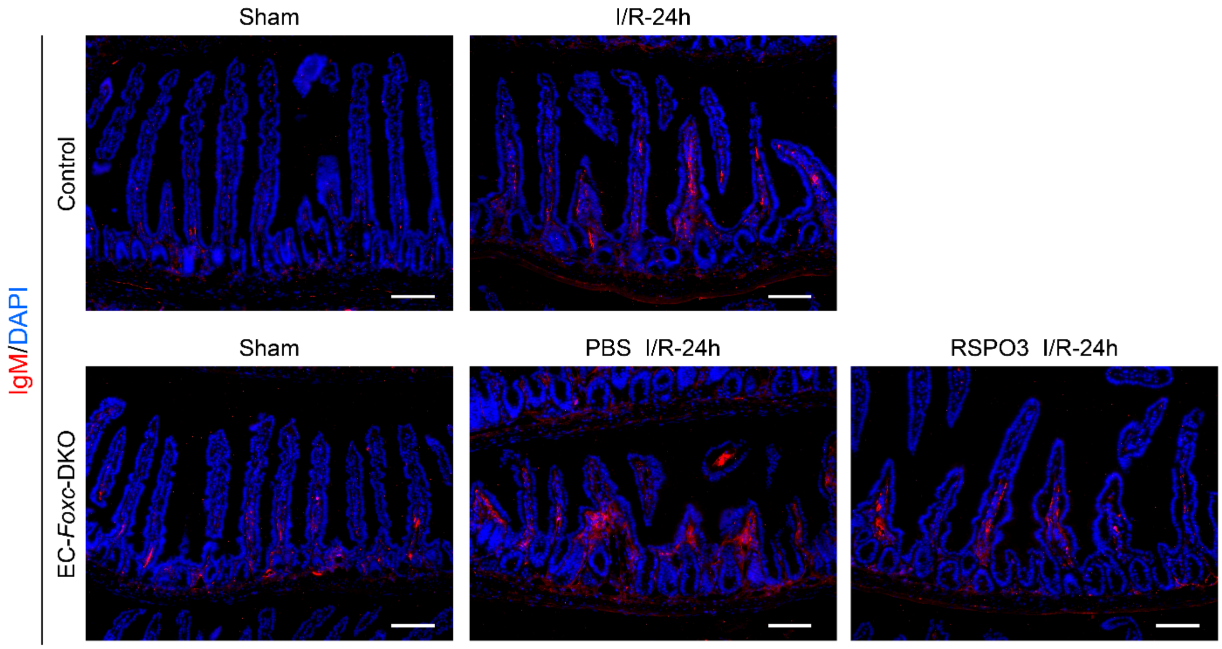


**Supplementary Figure 9. RSPO3 alleviates IgM accumulation in the intestinal mucosa after I/R in EC-*Foxc*-DKO mice.** Representative immunostaining images of small intestine labeled with IgM in control and EC-*Foxc*-DKO mice in sham or at 24h after I/R. After I/R, more IgM accumulation in the intestinal mucosa is found in EC-*Foxc*-DKO mice compared with control mice. The accumulation of IgM is alleviated in RSPO3-treated compared with PBS-treated EC-*Foxc*-DKO mice. Paraffin sections (4 μm), scale bars = 100 μm.

#### Supplementary Figure 10

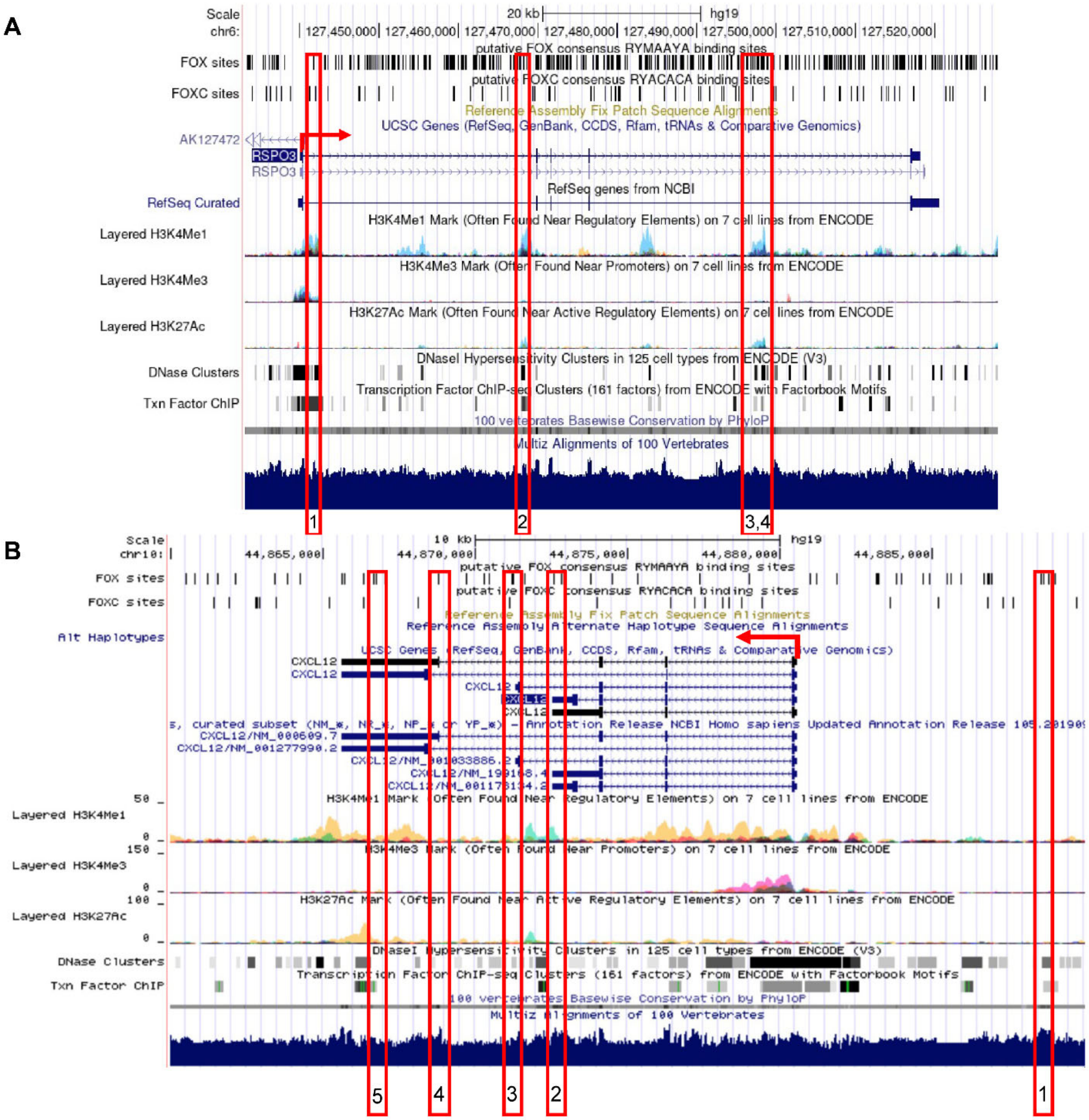


**Supplementary Figure 10. *In silico* identification of putative FOX-binding sites in the *RSPO3* and *CXCL12* loci.** Putative FOX-binding sites in regions of the human *RSPO3* **(A)** and *CXCL12* **(B)** loci as viewed on the UCSC genome browser (https://genome.ucsc.edu) [51]. Vertical lines on the 'FOX sites' and 'FOXC sites' tracks indicate putative binding sites corresponding to the FOX 'RYMAAYA' consensus sequence and the FOXC 'RYACACA' consensus sequence [70] predicted using the Hypergeometric Optimization of Motif EnRichment (HOMER) suite of tools [52]. Red boxes indicate evolutionary conserved regions (ECRs) containing FOX-binding sites between human and mouse genomes that are conserved and aligned as identified using the ECR browser tool (ecrbrowser.docde.org). The red arrows indicate the site of transcription initiation for *RSPO3* or *CXCL12*.

#### Supplementary Video Legends

**Supplementary Video 1. 3D structures of intestinal blood and lymphatic vasculatures in Control mice 18.5h after I/R**

**Supplementary Video 2. 3D structures of intestinal blood and lymphatic vasculatures in EC-*Foxc*-DKO mice 18.5h after I/R**

Representative 3D videos created using IMARIS software based on the confocal images of whole-mount immunostaining of distal jejunums show the 3D structures of intestinal blood capillaries (labeled with CD31, green) and lymphatic vessels (labeled with LYVE1, red) after I/R at 18.5h. In Control **(Supplementary Video 1)**, damages found at top of the villous blood vasculatures form openings to the capillary cage, accompanied by endothelial cell projections being formed for the repair. However, in EC-*Foxc*-DKO group **(Supplementary Video 2)**, the intestine loses most of its villi together with the villous blood and lymphatic vessels. The remaining villous, cryptal and submucosal vasculatures are also damaged and less branches can be found in the broken vessels. Lymphatic vessels are dilated in the cryptal area. Scale bars = 50 μm.

#### Supplementary Table 1

| Cluster | Marker | Cluster | Marker | Cluster | Marker | Cluster | Marker |
|---|---|---|---|---|---|---|---|
| BECs | Fabp4 | Enteroend-ocrine cell 2 | Rbp2 | Macrophage 1 | S100a9 | Plasmacyt-oid dendritic cells | Bst2 |
| | Ly6c1 | | S100a6 | | Cxcl2 | | Siglech |
| | Igfbp7 | | Lgals3 | | Il1b | | Ccl4 |
| | Plvap | | Tm4sf20 | | S100a8 | | Rnase6 |
| | Ly6a | | Serpinb6a | | G0s2 | | Mpeg1 |
| | Ptprb | | Anxa2 | | Srgn | | Lsp1 |
| | Flt1 | | Sprr2a3 | | Tyrobp | | Ly6c2 |
| | Pecam1 | | Gsta1 | | Cebpb | | Cybb |
| | Cd36 | | Fam162a | | Cd14 | | St8sia4 |
| | Epas1 | | Pmp22 | | Fcer1g | | Ccr9 |
| Bottom enterocytes | Reg3b | Erythroid cells | Hba-a1 | Macrophage 2 | Lyz2 | Telocytes /Trophocytes | Dcn |
| | Fabp1 | | Hbb-b1 | | Ccl6 | | Col3a1 |
| | Reg3g | | Hba-a2 | | Ccl2 | | Gsn |
| | Plac8 | | Alas2 | | Ccl9 | | Col1a2 |
| | Sis | | Ube2l6 | | Mafb | | Col1a1 |
| | Mgst1 | | Snca | | Ctsb | | Mgp |
| | Maoa | | Fech | | Psap | | Sfrp1 |
| | Arg2 | | Cyb561d1 | | Ctsc | | Lum |
| | Aldh1a1 | | Irgc1 | | Lgmn | | Bgn |
| | Gstm3 | | Nudt15 | | Ms4a6c | | Serping1 |
| CD4+ T cells | Emb | Fibroblasts | Apoe | Macrophage 3 | C1qa | Top Enterocytes | Apoa4 |
| | Lat | | Scn7a | | C1qb | | Ada |
| | Tcf7 | | Sparc | | C1qc | | Apob |
| | Ms4a4b | | Apod | | Acp5 | | Apoa1 |
| | Itgb7 | | Fxyd1 | | Apol7c | | Clca4a |
| | Cd69 | | Prnp | | Il22ra2 | | Apoc3 |
| | Lef1 | | Cryab | | Dnase1l3 | | 2010109I03Rik |
| | Bcl11b | | Plp1 | | Pla2g2d | | Selenop |
| | Cd27 | | Chl1 | | Batf3 | | Nt5e |
| | Arl4c | | Pmepa1 | | Tctex1d2 | | Dnpep |
| CD8+ T cells | Ccl5 | Follicular B cells | Cd74 | Middle Enterocytes | Fabp2 | Tuft and M cells | Krt18 |
| | Gzma | | H2-Ab1 | | Guca2b | | Cd24a |
| | Cd7 | | Ly6d | | Spink1 | | Adh1 |
| | Cd3g | | Vpreb3 | | Anpep | | Hck |
| | Rgs1 | | Bank1 | | Aldob | | Sh2d6 |
| | Nkg7 | | Cd83 | | Clca4b | | Lrmp |
| | Gzmb | | Pou2f2 | | Smim24 | | Selenom |
| | AW112010 | | Gpr183 | | Slc5a1 | | Tm4sf4 |
| | Cd8a | | H2-DMb2 | | Leap2 | | Dclk1 |
| | Cd3e | | Fcmr | | Guca2a | | Avil |
| Crypt cells | Krt19 | Goblet and Paneth cells | Zg16 | Myocytes /Pericytes | Acta2 | | |
| | Dmbt1 | | Tff3 | | Tagln | | |
| | Gpx2 | | Fcgbp | | Myh11 | | |
| | Hmgb2 | | Spink4 | | Sparcl1 | | |
| | H2afz | | Clca1 | | Tpm1 | | |
| | Birc5 | | Lgals2 | | Tpm2 | | |
| | Ube2c | | Agr2 | | Cald1 | | |
| | Cks2 | | Lypd8 | | Myl9 | | |
| | Pclaf | | Gm1123 | | Rgs5 | | |
| | Ccdc34 | | Ido1 | | Flna | | |
| Enteroend-ocrine cell 1 | Sct | LECs | Mmrn1 | Plasma cells | Jchain | | |
| | Chgb | | Ccl21a | | Mzb1 | | |
| | Chga | | Timp3 | | Pou2af1 | | |
| | Neurod1 | | Lyve1 | | Nap1l1 | | |
| | Cpe | | Cavin2 | | Sec11c | | |
| | Krt7 | | Reln | | Dut | | |
| | Fxyd3 | | Aqp1 | | Eaf2 | | |
| | Reg4 | | Flt4 | | Mef2b | | |
| | Tph1 | | Fgl2 | | H2afx | | |
| | Pcsk1 | | Tshz2 | | Top2a | | |

**Supplementary Table 1. Top 10 marker genes used for the identification of specific cell clusters in scRNA-seq data analysis**

#### Supplementary Table 2

| Primary Antibodies | | | | | |
|---|---|---|---|---|---|
| **Antibody** | **Supplier** | **Catalog number** | **Host Species** | **Clonality** | **Application** |
| β-actin | Sigma | A1978 | mouse | mAb | WB |
| β-catenin | Santa Cruz | sc-59737 | mouse | mAb | IHC-P |
| BrdU | Abcam | ab6326 | Rat | mAb | IHC-P |
| CCND1 | Invitrogen | MA5-14512 | Rabbit | mAb | IHC-P |
| CD31 | BD Biosciences | 553370 | Rat | mAb | WM, IHC-F |
| CD31 | Cell signaling | 77699 | Rabbit | mAb | IHC-P |
| CD31 | R&D | AF3628 | Goat | pAb | WM |
| EMCN | Abcam | ab106100 | Rat | mAb | IHC-P |
| EpCAM | Cell signaling | 93790S | Rabbit | mAb | IHC-P |
| FOXC1 | Abcam | ab227977 | Rabbit | mAb | WM |
| FOXC2 | Kind gift from Dr. N Miura (Miura et al., 1997, Genomics) | - | Rat | mAb | IHC-P |
| GFP | Invitrogen | A-11122 | Rabbit | pAb | IHC-F, WB |
| IgM | Invitrogen | 61-6800 | Rabbit | pAb | IHC-P, WB |
| LYVE-1 | Abcam | ab14917 | Rabbit | pAb | WM, IHC-P, IHC-F |
| LYVE-1 | R&D | AF2125 | Goat | pAb | WM, IHC-P, IHC-F |
| OLFM4 | Cell signaling | 39141T | Rabbit | mAb | IHC-P |
| PROX1 | R&D | AF2727 | Goat | pAb | IHC-P |
| RSPO3 | Sigma | HPA029957 | Rabbit | pAb | IHC-P |
| VEGFR2 | R&D | AF644 | Goat | pAb | WM |
| VEGFR3 | R&D | AF743 | Goat | pAb | WM |

| Secondary Antibodies | | | | |
|---|---|---|---|---|
| **Antibody** | **Reactivity** | **Host Species** | **Supplier** | **Application** |
| Alexa 405-conjugated | Rabbit | Donkey | Jackson Immuno Research Labs | WM, IHC-P, IHC-F |
| Alexa 488-conjugated | Rat/Rabbit/Goat | Donkey | Thermo Fisher | |
| Alexa 568-conjugated | Rat | Donkey | Abcam | |
| Alexa 568-conjugated | Rabbit/Goat | Donkey | Thermo Fisher | |
| Alexa 594-conjugated | Rabbit/Goat | Donkey | Thermo Fisher | |
| Alexa 594-conjugated | Mouse (IgG1) | Goat | Thermo Fisher | |
| Alexa 647-conjugated | Rat/Goat | Donkey | Thermo Fisher | |
| HRP conjugated | mouse | Goat | Thermo Fisher | WB |
| HRP conjugated | Rabbit | Goat | EMD | |

IHC-P: immunohistochemistry staining on paraffin sections; IHC-F: Immunohistochemistry staining on frozen sections; WB: Western Blot; WM: Whole-mount staining.

#### Supplementary Table 2. Antibodies used for whole-mount, sections and Western blot

**Supplementary Table 3**

| Gene | Forward | Reverse |
|---|---|---|
| *Cox-2* | GCATTCTTTGCCCAGCACTT | GGCGCAGTTTATGTTGTCTGT |
| *Cxcl-12* | GCTCTGCATCAGTGACGGTAA | CGTGCAACAATCTGAAGGGC |
| *Foxc1* | TTCTTGCGTTCAGAGACTCG | TCTTACAGGTGAGAGGCAAGG |
| *Foxc2* | AAAGCGCCCCTCTCTCAG | TCAAACTGAGCTGCGGATAA |
| *IL-6* | AGAGGATACCACTCCCAACAGA | CCACGATTTCCCAGAGAACA |
| *Rspo3* | TGTGAGGCCAGTGAATGGAG | ATCTCGGACCCGTGTTTCAG |
| *TNF-α* | ACAGAAAGCATGATCCGCGA | CTGCCACAAGCAGGAATGAG |
| *18S* | GAAACTGCGAATGGCTCATTAAA | CCACAGTTATCCAAGTAGGAGAGGA |

**Supplementary Table 3. Primers used for qPCR analysis**
